# A machine learning based method for the identification of functionally important genes

**DOI:** 10.1101/2022.06.08.495277

**Authors:** Gourab Das, Indira Ghosh

## Abstract

Deciphering complex mechanisms underlying communicable and non-communicable diseases require comprehensive analysis of genetic factors and interactions between them. Experimental identification of genes related to pathogen’s virulence and human diseases is effective but laborious, time consuming and costly. Consequently, many genetic factors associated with pathogenesis or disease still remain to be unknown. In present work, a normalized point wise mutual information (nPMI) driven computational method has been developed to identify the association between biological entities (genes) and physiological responses utilizing published literatures in PubMed. Association prediction models are then developed using machine learning approach for four different datasets including virulent genes from two bacterial species (*E. coli* and *S. enterica*) and stress responsive genes from two plant species (*A. thaliana* and *O. sativa*). This approach provides a generic platform for identifying association of genes in diverse biological systems (host and pathogen) and provide up-to-date estimate of association measures of the genes with stress, virulence. In future, the causal relation between them may be of research importance.

## Introduction

Bacterial pathogenesis and crop’s stress adaptation are the inevitable prime biological events with direct sociological impacts across globe [1-3]. These phenotypic consequences are driven by one or more genomic elements including their products, and underlying interactions among them [4-6]. In this context, detection of functionally important genes like virulence genes, stress responsive genes should be a mandate operation to perform. Among the experimental techniques, microarray, PCR and multiplex PCR based methods are notable but laborious, time and cost expensive jobs. Eventually, various computational methods have been developed to predict and characterize such important genes.

Homology based, machine learning based and network based approaches are the common existing virulent and stress responsive gene detection strategies [7-8]. Among them, traditional homology based methods finds genes with same functions and share high percentages of sequence similarities or identities. A web-server, VRprofile [9] uses such method for searching virulence, antibiotic resistance genes in newly sequenced chromosomes. But this method is often misleading for distant relatives having low sequence similarities. To overcome this limitation, network based approach has been adopted which predicts virulent factors using the protein-protein interaction network database STRING [10]. Though this method performs better than others but tragically underperform where network prediction is poor as in case of *Mycobacterium tuberculosis* H37Rv [7]. Moreover, lack of interactive interface restricts its usability. On the other hand, machine learning based tools like Virulent-GO [11], Virulent-Pred [12], VICMpred [13] are popular ones but MP3 [14] outperforms in prediction accuracy with the availability of both web-server and standalone package. MP3 provides a hybrid approach comprising of Support Vector Machine (SVM) and Hidden Markov Model (HMM) based methods to predict pathogenic proteins from genomic and metagenomic datasets. It uses di-peptide composition as features for building SVM models and Pfam domains for HMMs. But these kinds of machine learning based models often suffer from the problem of optimum feature selection and create a space for developing new methods with optimized feature sets. For example, MP3 is only intended for the prediction of pathogenic proteins; hence use of other features like DNA sequence, text, network, gene ontology (GO) is limited which might enhance the classification accuracy. Moreover, earlier studies has shown inclusion of GO based features along with sequence features improves model’s accuracy [11].

On the contrary, very few prediction algorithms are available for bacterial stress gene detection. Limited studies have been performed to distinguish stress-related genes in diverse bacterial species and strains. Earlier and recent reviews have documented the bacterial stress-responsive genes which remains the major primary resource of bacterial stress-related genes [15-17]. Among them, reviews on bacterial oxidative stress-related gene by Ezraty et al. [18], on the stress-responsive genes during host infection by Fang et al. [19], genes require for survival under competitive microbial society by Conforth et al. [20], generalized stress-related genes by Giuliodori et al. [21], Han et al. [22], Boor et al. [23], stress response leading to antimicrobial resistance by Fruci et al. [24] are highly recommendable. Moreover, a study on abundance and role of repetitive sequences in the bacterial stressed genes by Rocha et al. and others [25-26] are highly commendable. In the same context, rigorous experimental detection of stress-related genes in *Mycobacterium tuberculosis* by Schoolnik’s group is highly appreciable [27].

On the other hand, identification of plant stress-responsive genes is much more challenging because of complexity and larger genome size. But many efforts have been done to classify plant stress-related genes in major crop species like rice, wheat, barley, maize and model species like *Arabidopsis* etc. Several databases have been also developed to list plant stress-responsive genes including *Arabidopsis* stress gene database by Borkotoky et al. [28], plant stress gene catalogue by Wanchana et al. [29], Stress-responsive transcription factor database by Naika et al. [30], generalized stress genes for different plant species by Prabha et al. [31], Papdi et al. [32], resistance gene database by Sanseverino et al. [33], plant stress-related protein database (PSPDB) by Kumar et al. [34], drought stress related genes in DroughtDB by Alter et al. [35] etc. Majority of these databases have been identified genes by either microarray expression analysis or homology based methods. Also, computational methods for plant stress gene detection from expression data have been reviewed recently.

In the present work, a new text based feature called “Literature Association (LitAsso)” score has been introduced along with other DNA and protein sequence based features. Then, a machine learning based multi-model-method classification framework has been developed for comparing the many models developed applying many classification methods to distinguish the genes associated with virulence and stress adaptation. A web-server has been also employed for quick and automated computational identification of these functionally important genes. It takes minimal input of gene symbol(s) from user and predicts its class (positive/negative). These lists of predicted genes can be opted for further experimental validation to unwrap complex mechanisms behind these phenotypic responses.

## Materials & Methods

To date several binary classification methods are available including logistic regression (LR), linear discriminant analysis (LDA), SVM, random forests, artificial neural networks (ANN), nearest neighbour and many more. Each method classifies the positive (here, virulent and stress responsive gene) from negative (housekeeping genes) sets on the basis of learning from some training data and thresholds. To build multi-model-method framework many LR, LDA, and SVM models have been developed and compared to find the best one in terms of accuracy. Many sequence based features have been calculated from the sequences available in the published databases like NCBI nucleotide, Virulence Factor database (VFDB) and used to train the models. Along with these well-known sequence based features, a new text based feature based on literature association (LitAsso) has been implemented by calculating normalized pointwise mutual information (nPMI) score [36]. Generally, to find the correlation/association between two variables, several statistical techniques and formulations have been utilized which includes Pearson, Spearman, Kendall, Matthews’s correlation coefficient, Association rule mining (ARM) [37]. nPMI is preferred over aforementioned correlation metrics as it requires only the prior knowledge on distribution function not the type of underlying relations e.g. linear, non-linear, monotonic etc. [38]. The classification models have been trained on a part of published data and tested on the rest part as blind test set which has not been used during training. A workflow of the developed method has been depicted in the Figure 1.

**Figure 1:**
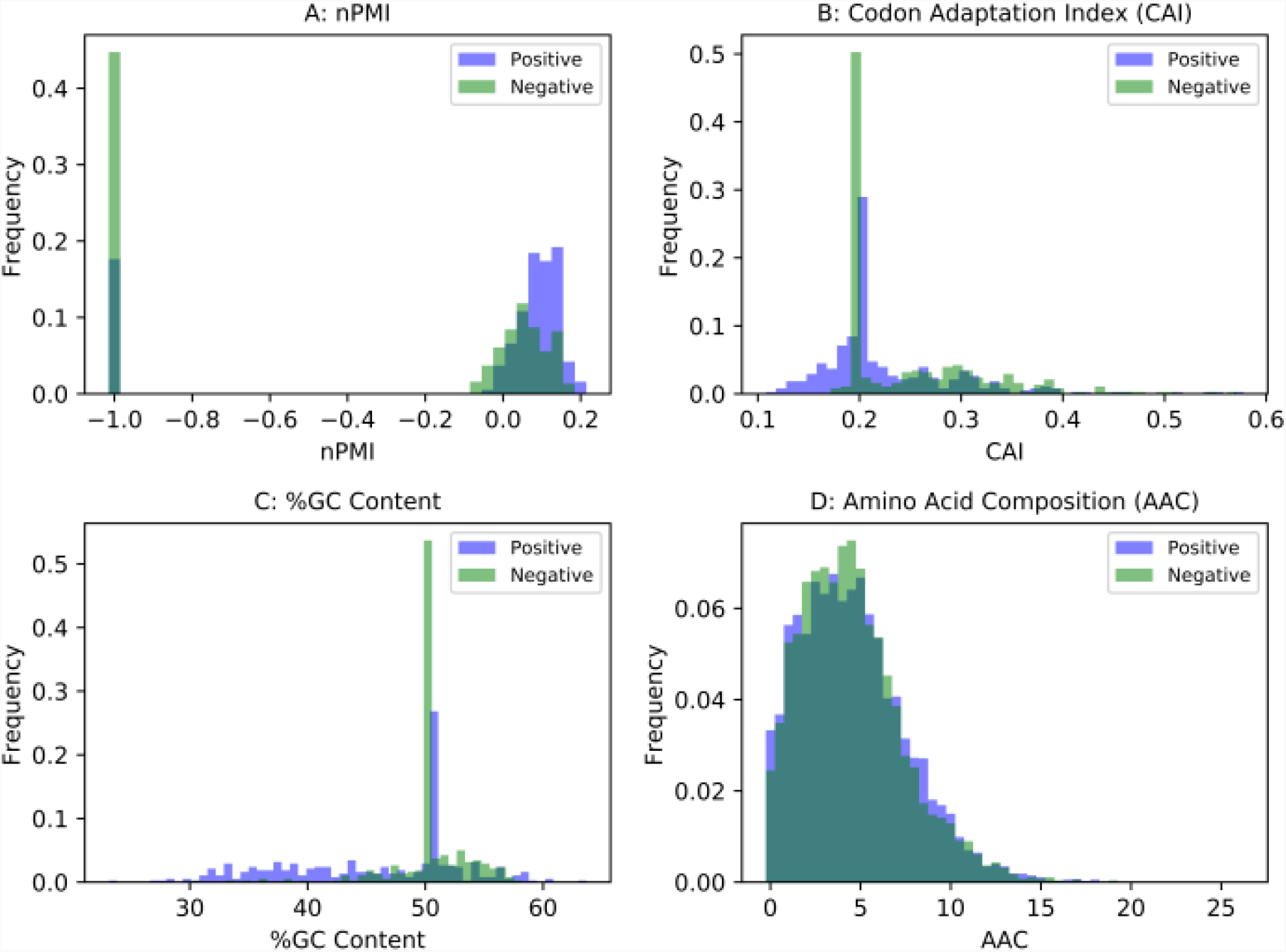
Distribution of features in the *E. coli* virulence related dataset. A: Normalized Pointwise mutual information (nPMI); B: Codon adaptation index (CAI); C: %GC Content; D: Amino acid composition (AAC); Blue: Positive set; Green: Negative set.

### Datasets

So far, many public databases are available comprising of bacterial virulence genes which are either identified by experimental procedures or collected from published literatures. Compiling these databases namely VFDB [39], Victors database from Pathogen-Host Interaction Data Integration and Analysis System (PHIDIAS) [40] and Pathosystems Resource Integration Center’s (PATRIC) [41] virulence factors data, a list of total 445 unique symbols of *E. coli* virulent genes have been prepared. This set is called as positive virulence dataset (Supplementary Table T1). To check the robustness of the method, another class of functionally important genes i.e. stress responsive genes from higher biological systems like plants have been selected. This set of positive data contains 413 *Arabidopsis* stress responsive genes which has been collected from published literatures [42] and named as positive stress data (Supplementary Table T2). The stressors include abiotic such as water deficit (drought), salinity, excess metal and biotic factors like nematode stress. *E. coli* and *Arabidopsis* housekeeping (HK) gene symbols with DPM^1^ values closer to zero have been extracted from PaGenBase database and utilized as negative virulence and stress responsive sets respectively (Supplementary Table T3 and T4) [43]. To check the consistency in prediction performance, two additional positive datasets i.e. 336 *Salmonella enterica* virulent genes (Supplementary Table T5) from aforementioned bacterial virulent gene databases and 900 stress responsive genes from *Oryza sativa* (Supplementary Table T6) have been also included from published literatures [30-31, 35, 44-46]. Common gene symbols between *E. coli* negative set and *S. enterica* genes have been used to build the negative dataset for *S. enterica*. Finally, excluding the symbols which are available in *S. enterica* positive virulence dataset, *S. enterica* negative virulence gene list has been prepared (Supplementary Table T7). For *O. sativa*, HK genes from Rice Expression Database (RED) [47] have been chosen to build the negative stress responsive gene set for rice (Supplementary Table T8). Following the authors, selection of the HK genes has been done using the tissue specificity index (τ < 0.95).

### Calculation of text based feature LitAsso using PubMed

Set of virulent, and stress related keywords and phrases are selected from knowledgebase and used to build PubMed advanced queries (Supplementary Table T9). While publishing any literature that reports genes related to stress, virulence or any disease, it is highly probable that authors will mention keywords like ‘stress’, ‘resistance’, ‘virulence’ or ‘pathogenesis’ etc. in vicinity of gene symbols in context to express its functional implications if any exists. Using this generality in art of writing, nPMI scores have been calculated by checking co-occurrences of a gene symbol and set of keywords and phrases in PubMed abstracts. Biopython (v1.69) modules for NCBI E-utilities have been applied for this calculation. For each gene symbol, nPMI score for last past 28 years (1990 to 2017) is used as the LitAsso feature. This particular time interval is used as before this interval, very less number of publications are found related to stress, virulence or diabetes and not sufficient for estimating nPMI values.

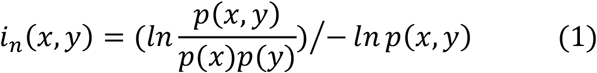

where, *x* is gene symbol, *y* is set of keyword(s) and phrases combined with boolean operators like OR, AND etc. *p(x), p(y)* are the probability of finding *x, y* and *p(x, y)* is the joint probability of *x* and *y*.

### Sequence based features

Apart from text based feature, many DNA and amino acid sequence based features have been selected under present study. Among the DNA sequence-based features, %GC content and Codon adaptation index [48] have been preferred because earlier reviews [49] have shown that %GC content does influences stress adaptation in bacteria. However, the role of %GC content in plants has not been clearly understood but its association with genomic adaptation leading to evolution has been explored [50]. Protein sequence-based feature include amino acid composition (AAC) which is the relative abundance of each amino acid in a particular protein sequence [51]. DNA sequence features have been calculated using Biopython and amino acid features using pydpi package (v1.0).

### Feature selection

Therefore, total 23 features including 2 DNA sequence feature, 20 protein sequence features and 1 text based feature have been utilized for classifier training. Prior to training, to check the influences of these parameters on classifier’s performance, principal component analysis (PCA) has been performed for each datasets to select the optimum set of features. As some features can have zero values, log transformation would not be a good option to select instead Singular value decomposition (SVD) has been used as it is more efficient and robust than other transformation approaches. PCA has been performed using standard R (v3.4) base packages.

### Selection of blind test dataset

To avoid any kind of bias during classifier learning due to class imbalance, positive to negative class ratio has been maintained as ∼1:1 in the positive-negative combined datasets for each species. It is hard to make an assumption on the ideal train-test split ratio for any classifier design but should be chosen carefully as it influences prediction performance [52]. So, for each species, combined dataset is divided randomly into training and blind test sets with different proportions ranging from 50:50 to 90:10 preserving 1:1 positive to negative ratio. Any sample from this blind test set has not been used in classifier training to achieve an unbiased prediction. For cross-species analysis separate protocol has been followed for blind test set preparation. For phenotypic class plant stress, for each species blind test has been prepared by selecting 10% of the data after excluding the inter-species common symbols and preserving 1:1 positive to negative ratio in the blind test set. But in case of bacterial virulence since negative sets are common, after excluding the inter-species common symbols, 10% and 20% samples have been chosen from E. coli and S. enterica positive stress datasets respectively. After that training sets have been manipulated to preserve the 1:1 positive to negative ratio.

### Building multi-model-method framework

Three different methods have been chosen which include two linear i.e. linear discriminant analysis (LDA) and logistic regression and one non-linear supervised classification scheme. Support Vector Machine (SVM) has been opted as a non-linear classification scheme for building model and prediction into output classes (Positive/Negative). SVM with Radial basis function (RBF) kernel has been used along with tuning of cost (C) and gamma parameter. The gamma parameter defines how far the influence of a single training example reaches, with low values meaning ‘far’ and high values meaning ‘close’. The gamma parameters can be seen as the inverse of the radius of influence of samples selected by the model as support vectors. The C parameter trades off misclassification of training examples against the simplicity of the decision surface. A low C makes the decision surface smooth, while a high C aims at classifying all training examples correctly by giving the model freedom to select more samples as support vectors. To build this multi-model-method framework, In SVM Cost (C) and gamma tuning, C values from 1-100 and gamma values from 10-6 to 10-1 have been scanned for choosing best-performing thresholds for these parameters. To reduce the over-fitting, K-fold repeated (10 times) cross-validation has been performed. For datasets with imbalanced positive-negative class ratio, Synthetic Minority Over-sampling Technique (SMOTE) [53] has been used during the training procedure.

### Evaluation metrics

For assessing the predictive capability of the classifiers, receiver operating characteristic curve (ROC) and Precision-Recall (PR) curve have been plotted. ROC can be generated by plotting true positive rate (probability of correctly detecting the positives as positives) (TPR or sensitivity) and false positive rates (probability of wrongly detecting negatives as positives). On the other hand, PR is the representation of sensitivity values upon positive predictive values. PR curve is more accurate than popularly used ROC plot when the class imbalance is present in the data [54]. Apart from that other metrics like F1-score, logLoss, Kappa etc. have been also calculated to achieve a consensus about the predictive ability of the models.

### Results & Discussion

While calculating the features, it has been found that 85%, 65%, 31% and 10.7%, of the gene symbols have valid values for all of the features in *E. coli, S. enterica, A. thaliana* and *O. sativa* datasets respectively (Supplementary Table T10, T11, T12 and T13). Loss of samples from the datasets is due to many reasons. First, lack of gene symbol-sequence mapping; Second, presence of nucleic and amino acid additional letter codes in the sequences which are not readable by the standard programs for CAI and AAC calculations; Third, presence of ‘N’ letter in the DNA sequences which causes the faulty codons resulting data loss. Basically, ‘N’ stands for many cases like gaps, sequencing error, instrument error etc. [55] which can cause faulty results. Hence, for achieving more accurate prediction, samples having all feature values have opted for classifier designing.

### Feature distributions

In the *E. coli* virulent dataset, differences in central tendency values (mean/median) between the positive and negative groups have been observed for rest of the features except %GC content and AAC. nPMI mean value for the positive set of genes is -0.08 with a variance of 0.18 whereas that of the negative set is -0.4 with a variance of 0.29 (Figure 1A) (nPMI range: -1 to +1). Little difference in CAI values between the positive (mean 0.23 with variance 0.004) and the negative group (mean 0.31 with a variance of 0.005) has been found to exist (Figure 1B). Like CAI feature, a little difference in median values (positive 49% and negative 50%) has been noticed between the groups in case of %GC content (Figure 1C). But, for both positive and the negative group, AAC median values are nearly equal to 4.5 (Figure 1D). In case of *S. enterica* dataset, differences in central tendency values have been observed in case of nPMI and %GC content only (Supplementary Figure F2). These results indicate that except AAC, other features have some ability to discriminate virulent and non-virulent genes in bacteria.

In case of *A. thaliana* stress dataset, only mean values of nPMI is found be little different between the groups. Mean value of nPMI in the positive group is -0.34 compared to that negative set which is -0.50 (Supplementary Figure F3A) is very distributed. Like other two datasets, variances within the groups are very high i.e. 0.28 for both positive and negative groups. CAI values between the groups (0.17 for positive and 0.18 for negative) are quite comparable having very less variances of 0.0004 (Supplementary Figure F3B). %GC content AAC are equal for in both groups and the median is nearly 38% and 4.7 respectively (Supplementary Figure F3C-F3D).

For, *O. sativa* dataset, mean nPMI and median %GC content values are found to be different between the groups (Figure F4). For all the datasets, only nPMI is found to be significantly different between the positive and negative groups but with high variances. Though, CAI feature has very less variance but it is not significantly different between the groups. %GC content and AAC have high variances and almost equal between the groups. These observations emphasize the importance nPMI feature for classifier design and novel gene prediction.

### Feature Selection using Principal Component Analysis (PCA)

PCA has been done with 23 features mentioned above to find the principal components and its loading values. For all the datasets 18-19 Principal Components (PCs) are needed to explain 80% and 95% of the variance of data (Figure 2A-2B).

**Figure 2:**
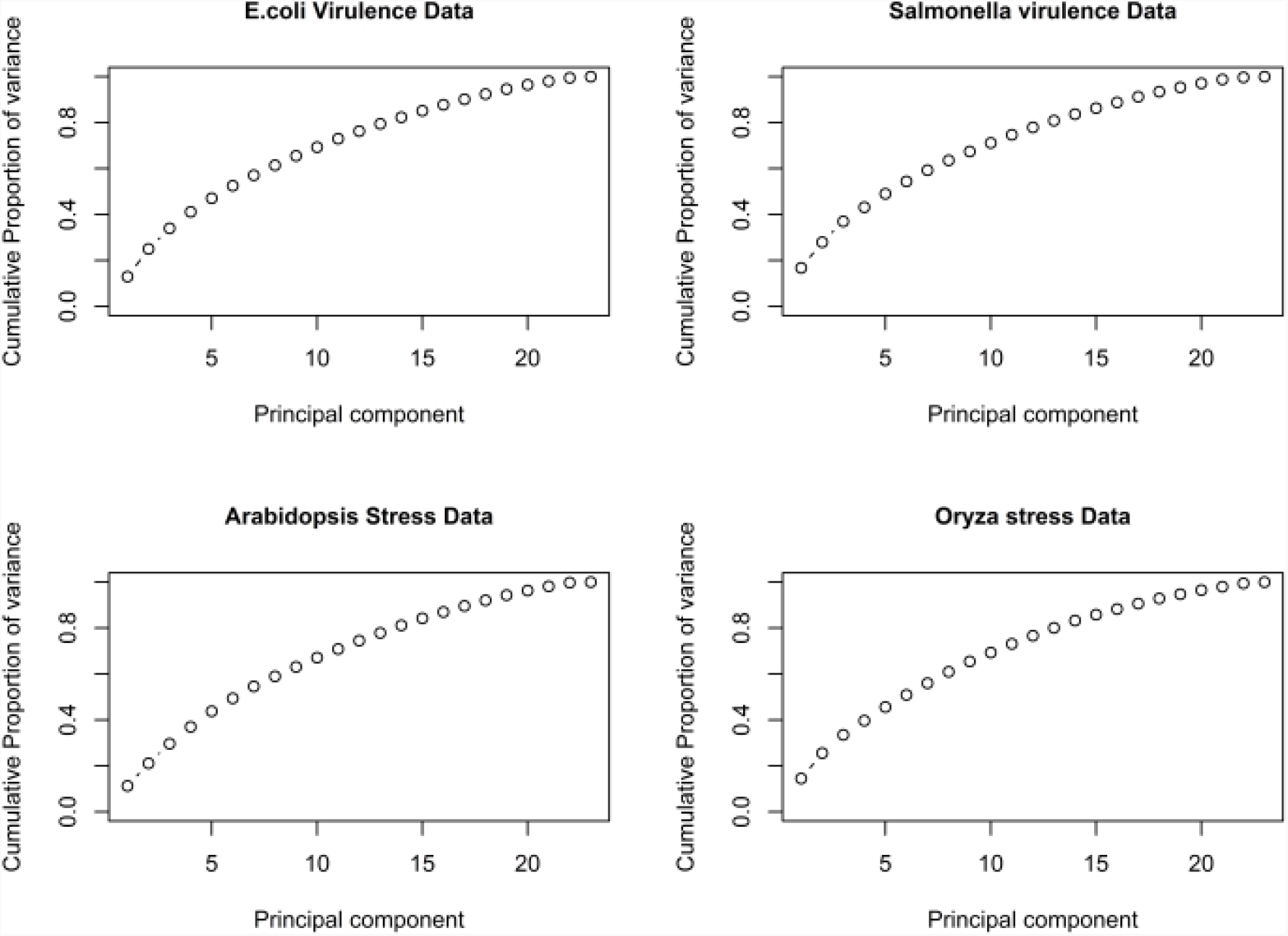
Feature selection using PCA. Cumulative proportion variances have been shown for PCs for all four datasets.

For *E. coli* virulence dataset nPMI (0.12), Asp (0.03), Phe (0.14), Ile (0.23), Lys (0.26), Asn (0.4), Ser (0.27), Thr (0.1) are the principal components whereas for *A. thaliana* stress data, nPMI (0.16), Ala (0.2), Cys (0.15), Gly (0.11), Phe (0.18), Ile (0.37), Met (0.034), Leu (0.29), Trp (0.15), Val (0.39) are found to be the important features. For contribution of each PC, variance has been plotted for both dataset in Supplementary Figure F5.

### Training and testing results

As mentioned earlier, three different classifiers have been used for training and testing. Several train-test ratios have been opted to split the data for learning and classification. Table 1 shows the summary of comparison of performances of best classifiers in terms of many evaluation metrics for all the datasets. Among them, as expected SVM is performing better than the others for all the datasets.

**Table 1:** Evaluation metrics summary of best models for all datasets

| Parameter and Metrics | *E. coli* virulence | *S. enterica* virulence | *A. thaliana* stress | *O. sativa* stress |
| --- | --- | --- | --- | --- |
| Samples (#) | 760 | 300 | 250 | 197 |
| Train : Test | 90:10 | 90:10 | 70:30 | 90:10 |
| Repeated CV* folds (#) | 10 | 5 | 3 | 3 |
| Best Model | Support vector machine (SVM) |  |  |  |
| Accuracy | 0.78 | 0.77 | 0.63 | 0.61 |
| Specificity | 0.73 | 0.81 | 0.53 | 0.50 |
| Sensitivity | 0.83 | 0.73 | 0.73 | 0.72 |
| Precision | 0.76 | 0.81 | 0.61 | 0.61 |
| F1 | 0.79 | 0.76 | 0.66 | 0.66 |
| AUC* | 0.86 | 0.84 | 0.65 | 0.64 |

\*CV: Cross-validation; AUC: Area under curve (ROC curve)

For bacterial virulence datasets, test accuracies for both the species are nearly equal however for plant stress datasets, it is very poor. SVM is performing better than other classifiers at a train-test cut-off ratio of 90:10. Utilizing only 23 features, this bacterial virulent gene classifier produces 78% accuracy which is almost quite less than that of current best tool for virulent protein prediction i.e. MP3 which uses many more features (approximately 50 times more in number) for classification with an accuracy of 89% for the genomic datasets and 96% for the proteomic datasets. As only 33% of the gene symbols have protein annotation in RefSeq indicating the requirement of virulent gene prediction tools at the genomic level where features of the predicted proteins only can be utilized. Distributions of evaluation metrics for *E. coli* dataset at train-test splitting ratio 0.9 has been presented in Figure 3. For, other datasets, evaluation metrics have been included in supplementary figures and tables (see readme.txt for description).

**Figure 3:**
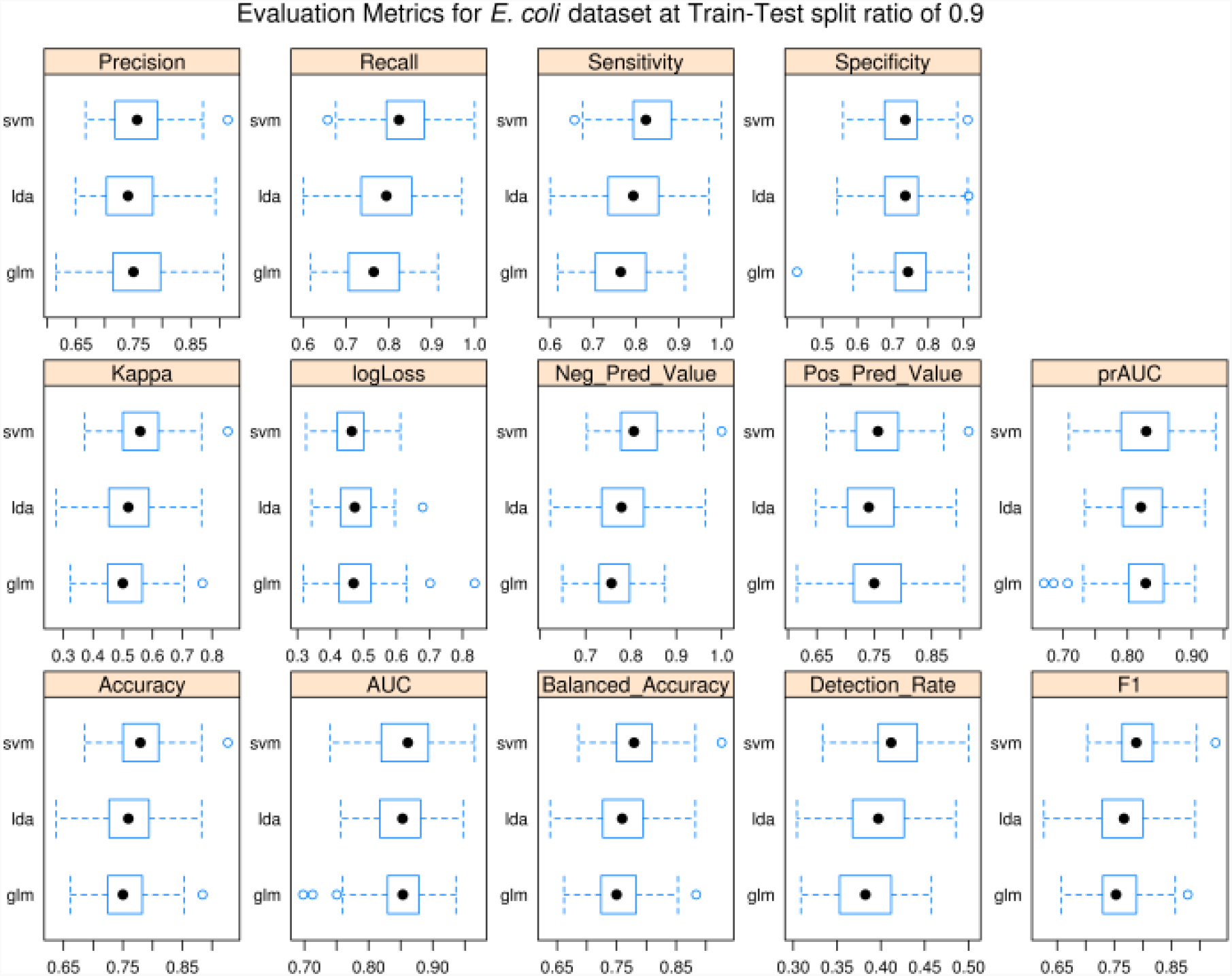
Distribution of evaluation metrics for *E. coli* dataset at train-test ratio 0.9

### Evaluation using ROC-PR Curve analysis

Concluding the model performance on the basis of cross-validation results solely would not be a wise idea to follow instead use of methods like ROC and PR analysis would be beneficial to perform a comprehensive and critical assessment of the model’s performance. Receiver operating characteristic or ROC curve analysis is one of the powerful tool for evaluating the performance of a binary classification model. It plots true positive rate vs. false positive rate to calculate the area under curve (AUC) score which in turn depicts the model’s performance. On the other hand, class imbalance (unequal numbers of positive and negative samples in the data) is a very usual and regular problem of machine learning. ROC curve is sensitive to the imbalanced dataset and can be misleading. Hence, Precision-Recall (PR) curve is required to be plotted for correct evaluations of the models. In the present study, plant stress is highly imbalanced, therefore; both ROC and PR curve analysis have been carried out in order to evaluate the classifiers performances. Performances of the classifiers trained with 23 features have been examined also using ROC and PR curve analysis^2^. While classifying Stress/Virulence genes, True positives (TP) are the positive samples (experimentally verified/literature driven stress/virulence related genes) which are predicted correctly whereas True negatives (TN) are the negative samples (housekeeping genes as predicted in PaGenBase database) which are also predicted correctly. False positives (FP) are the negative samples wrongly predicted as positives and False negatives (FN) are the positive samples wrongly predicted as negatives. FP initiates Type I error whereas FN introduces Type-II error. While comparing linear (LDA, LogReg) vs. non-linear (SVM with RBF kernel) classifiers, SVM performs little better than the linear models in case of all the datasets. For each dataset, classifiers producing best AUC values from ROC and PR curves have been plotted as shown in Figure 4 and Figure 5. According to the ROC and PR curves, SVM classifier’s performances are better than the other for all datasets (Figure 4 and 5). Fluctuations in the ROC and PR curves for plant stress datasets indicate classifier’s instability. It is also expected because of small training sample sizes (as mentioned in Results 3.1 section).

**Figure 4:**
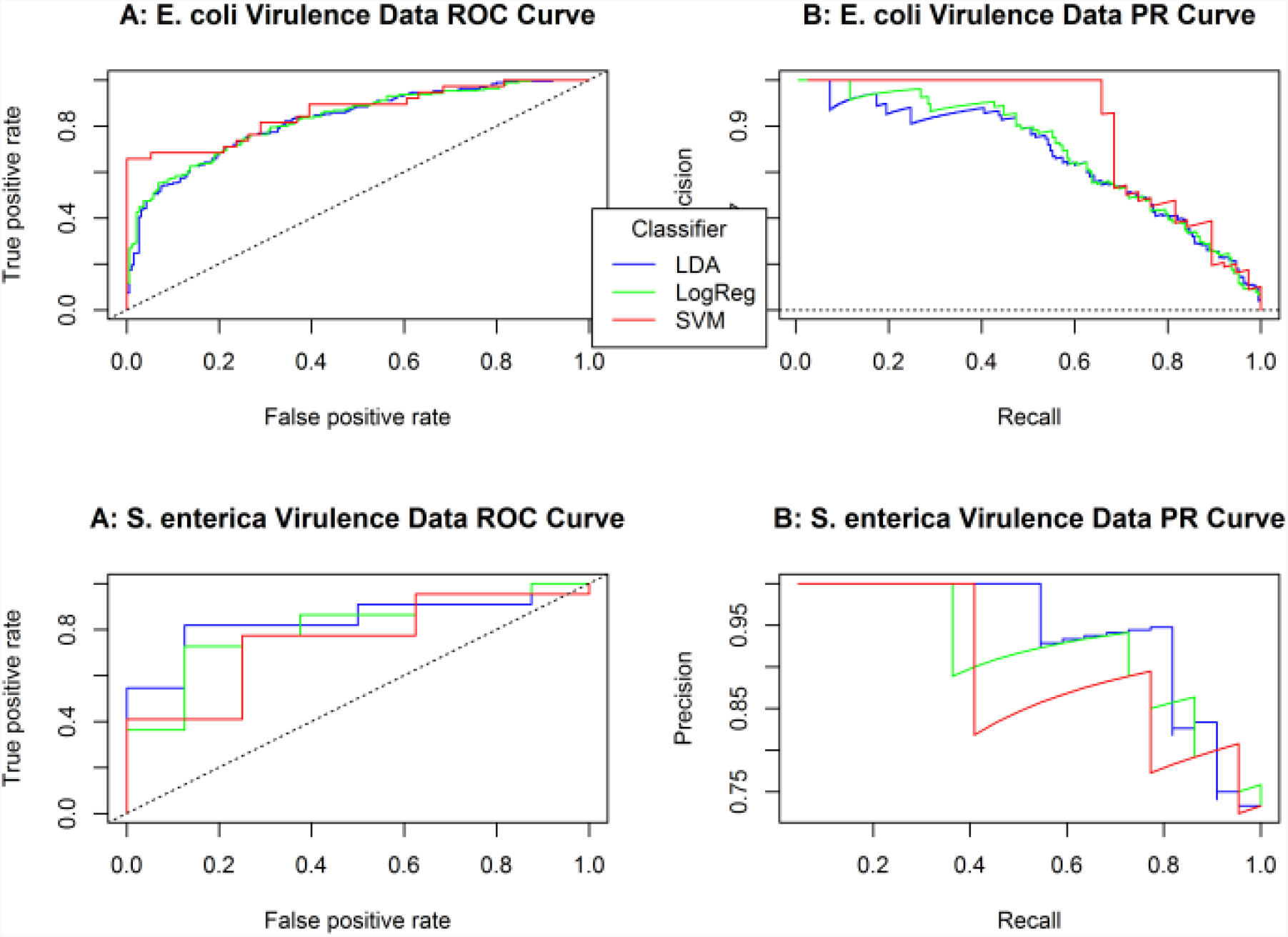
ROC and PR plot for *E. coli* and *S. enterica* virulence data sets; ROC: Receiver Operating Characteristics; PR: Precision-Recall; A: ROC curve for *E. coli* virulence dataset; B: PR curve for *E. coli* virulence dataset; C: ROC curve for *S*.*enterica* virulence dataset; D: PR curve for *S. enterica* virulence dataset; Blue: LDA; Green: Logistic Regression (LogReg); Red: SVM

**Figure 5:**
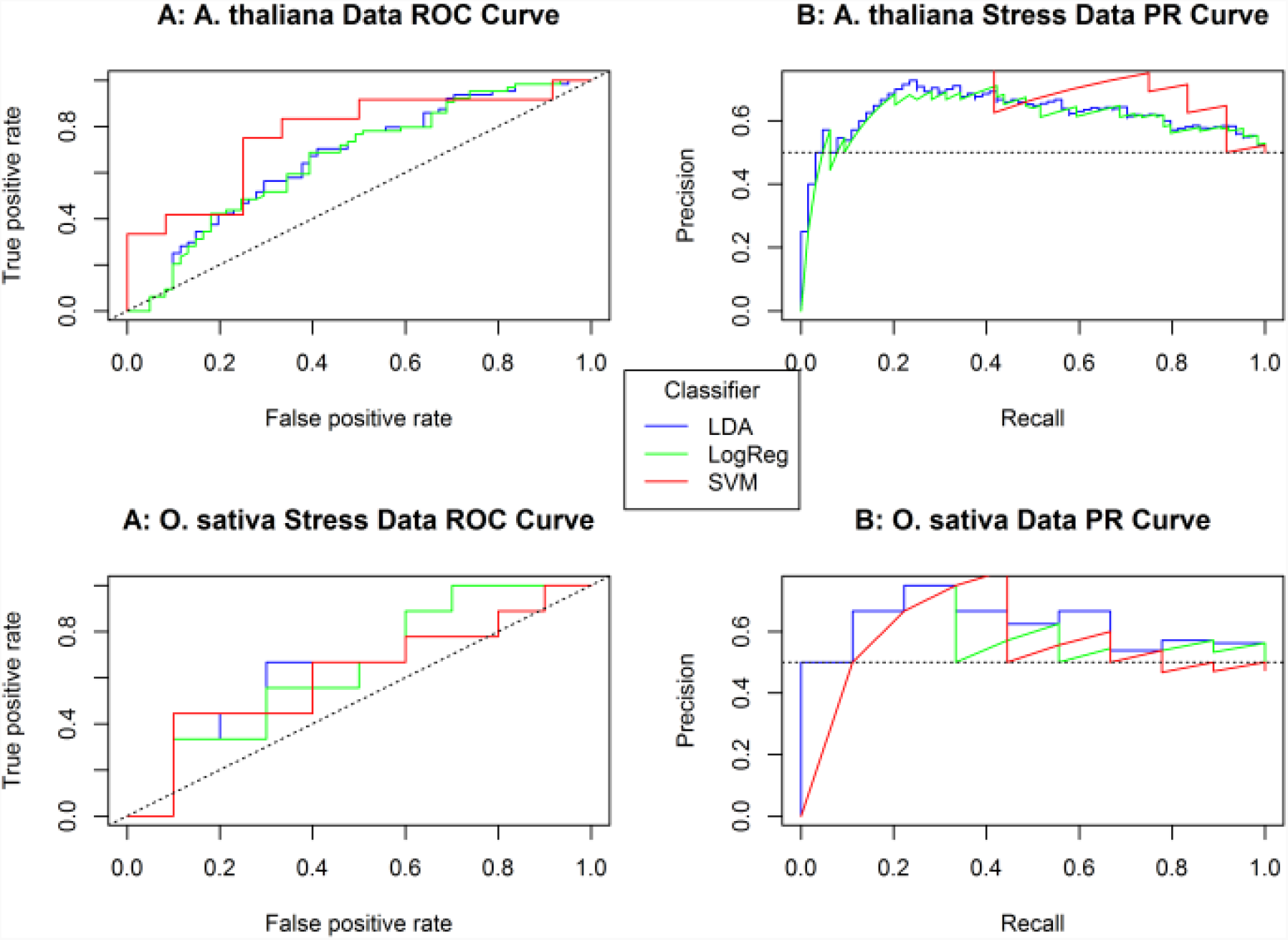
ROC and PR plot for plant stress data sets; ROC: Receiver Operating Characteristics; PR: Precision-Recall; A: ROC curve for *A. thaliana* dataset; B: PR curve for *A. thaliana* dataset; C: ROC curve for *O. sativa* dataset; D: PR curve for *O. sativa* dataset

### Cross-Species analysis

While summarizing the classifier’s performances from three different evaluation procedures i.e. ROC and PR curve analysis and cross-validation, consistent evaluations have been obtained from these different approaches. Bacterial virulence elated classification models are stable and performing better than plant stress classification models. The possible reasons are the insufficient data samples in the training and test sets, imbalanced positive (stress/virulence) and negative classes and continuous up-gradation of the actual class of the samples in the training data.

On checking the classifier’s performance using cross-species analysis, it has been observed that all of the classifiers have performed poorly. This indicates poor ability for generalization of the classifiers. Classifiers trained on the *E. coli* data can predict unknown *E. coli* genes efficiently (high test accuracy, AUC values as shown in virulent CV table and ROC plot for virulence data). But classifying *Salmonella* genes, classifiers have shown poor performance. The justification for this anomaly can be from the biological point of view. *E. coli* and *Salmonella* both are gram-negative Enterobacteriaceae and also phylogenetically very close to each other. But modes of pathogenesis, diseases caused are highly different. Similar results have been obtained in case plant stress datasets.

Earlier literature [55-58] have shown the high similarity in their core genomes i.e. a similar set of genes for executing basic cellular processes but significant differences in virulent gene profiles. Hence, it would be wise to build a separate classifier for each of the genus. For *Salmonella* and other genus, experimentally verified virulent gene symbols with DNA and protein sequences are also available in the VFDB data. Utilizing the proposed method classifiers can be constructed and evaluated. To obtain negative genes for virulence and stress for species other than *E. coli*, ortholog mining with a high sequence similarity can fulfill the requirement.

## Conclusions

Advancement in the sequencing technologies has created a big sequence data platform for studying and analysing genes and genomes related to disease, pathogenesis and adaptation under diverse stressful environments. But understanding causal relation between genotypic inputs and phenotypic outcomes require prior identification of associated genes. Experimental identification of these genes is a rigorous, laborious and time taking process. In this regard, the requirement of computational tools can be understood. In the present work, sequence and text features have been used together to train two linear (LDA and Logistic regression) and one non-linear (SVM) binary classifiers for 4 different gene datasets i.e. two bacterial virulence (*E. coli* and *S. enterica*), and two plant stress responsive gene sets (*A. thaliana, O. sativa*). DNA sequence-based features include codon adaptation index (CAI), %GC content whereas amino acid composition (AAC) has been used as the sole amino acid-based feature. A new feature i.e. nPMI association score derived from PubMed literature has been introduced which enhances the prediction ability of the sequence feature based classification models. While checking the influence of the individual feature on the data, it has been revealed that majority of the features or principal components are required to explain the underlying variance of the data (12 and 19 PCs out of 23 features can explain 80% and 90% of the variance). Therefore, all 23 features have been used for classifier training and performances have been evaluated by means of several methods including cross-validation, ROC-PR curve analysis, and re-sampling technique. Both linear and non-linear classifier’s performances are comparable but unavailability of the proper negative datasets and small sample size are the major issue behind the poor performances of *S. enterica* and plant classification models.

## Acknowledgements

GD is acknowledging Prof. Rocha

## Funding

This work has not been supported by any funding agency. But GD was supported by Council of Scientific and Industrial Research (CSIR) has provided Direct Senior Research Fellowship (SRF).

## Conflict of Interest

None declared.

## Footnotes

1 In PaGenBase database, DPM stands for dispersion measure which is calculated by mapping standard deviation of a gene expression profile between 0 and 1. Genes having DPM values < 0.3 are considered as housekeeping genes [43].

2 ROC and PR curves are the representations of trade-offs among the parameters 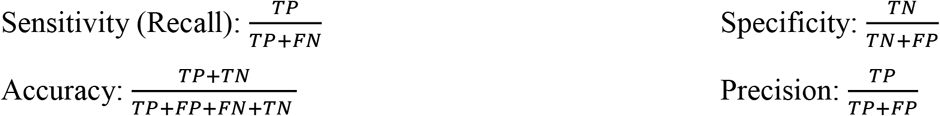

## Notes

### Competing Interest Statement

The authors have declared no competing interest.

