## supplementary_08062022 for "A machine learning based method for the identification of functionally important genes": readme.txt

ReadMe for nPMI Paper supplementary files

Tables Folder: supplementary_tables.xlsx: All the gene symbols (T1-T8), PubMed Advanced queries (T9);
Tables/feature_vectors: All the feature vectors for 4 datasets

==================================================

Figures Folder: All the histogram plots (F2-F4), PCA variance (F5)
Figures/CV-figures: Evaluation metrics plots for testset predictions for all the datasets at train-test split ratio ranging from 0.5 to 0.9


==================================================

Cross-validation-statistics Folder: Evaluation metrics statistics summary for testset predictions for all the datasets at train-test split ratio ranging from 0.5 to 0.9
