## Supplementary figures and images for "A machine learning based method for the identification of functionally important genes"

### F2.png

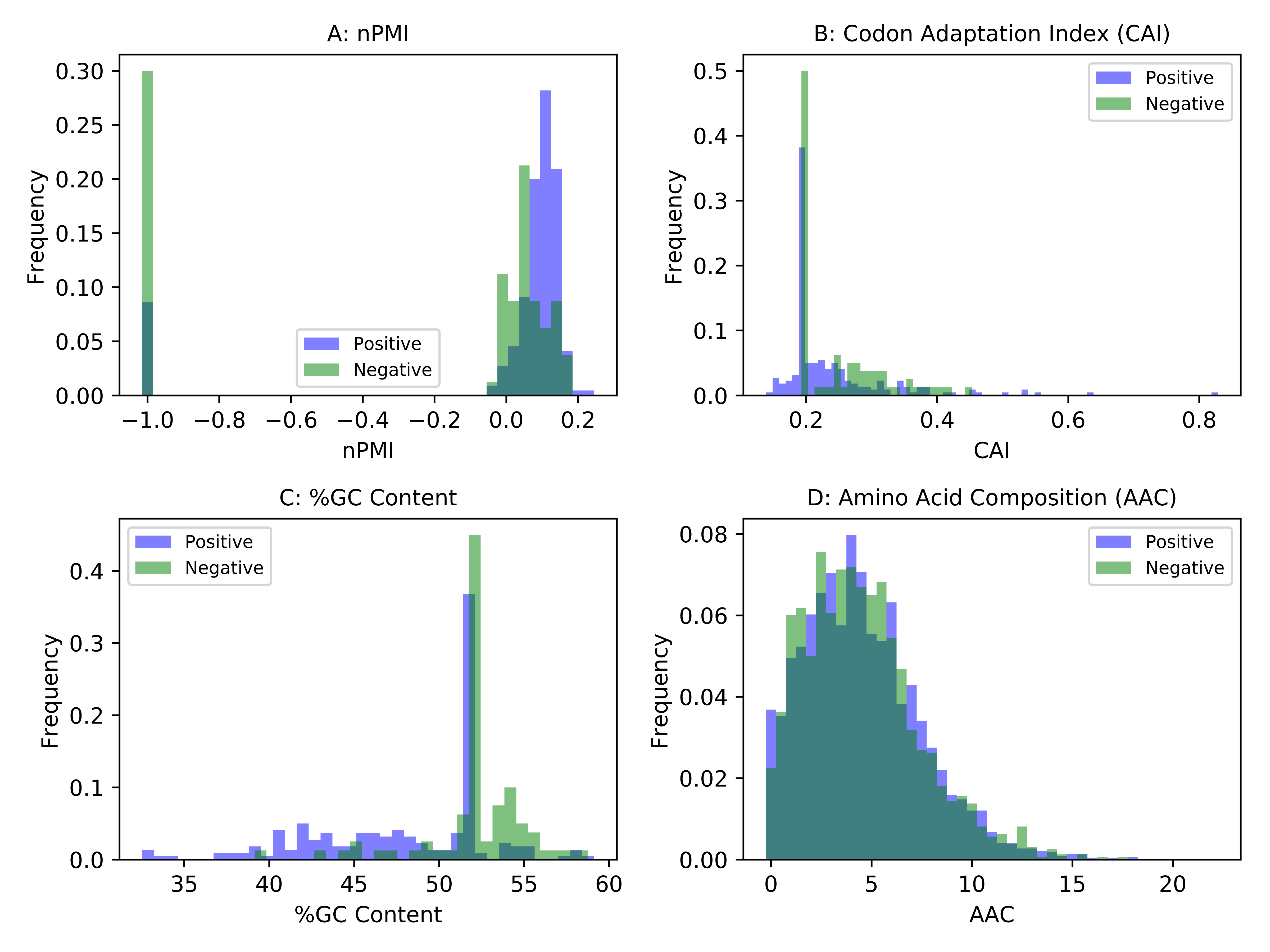

### F3.png

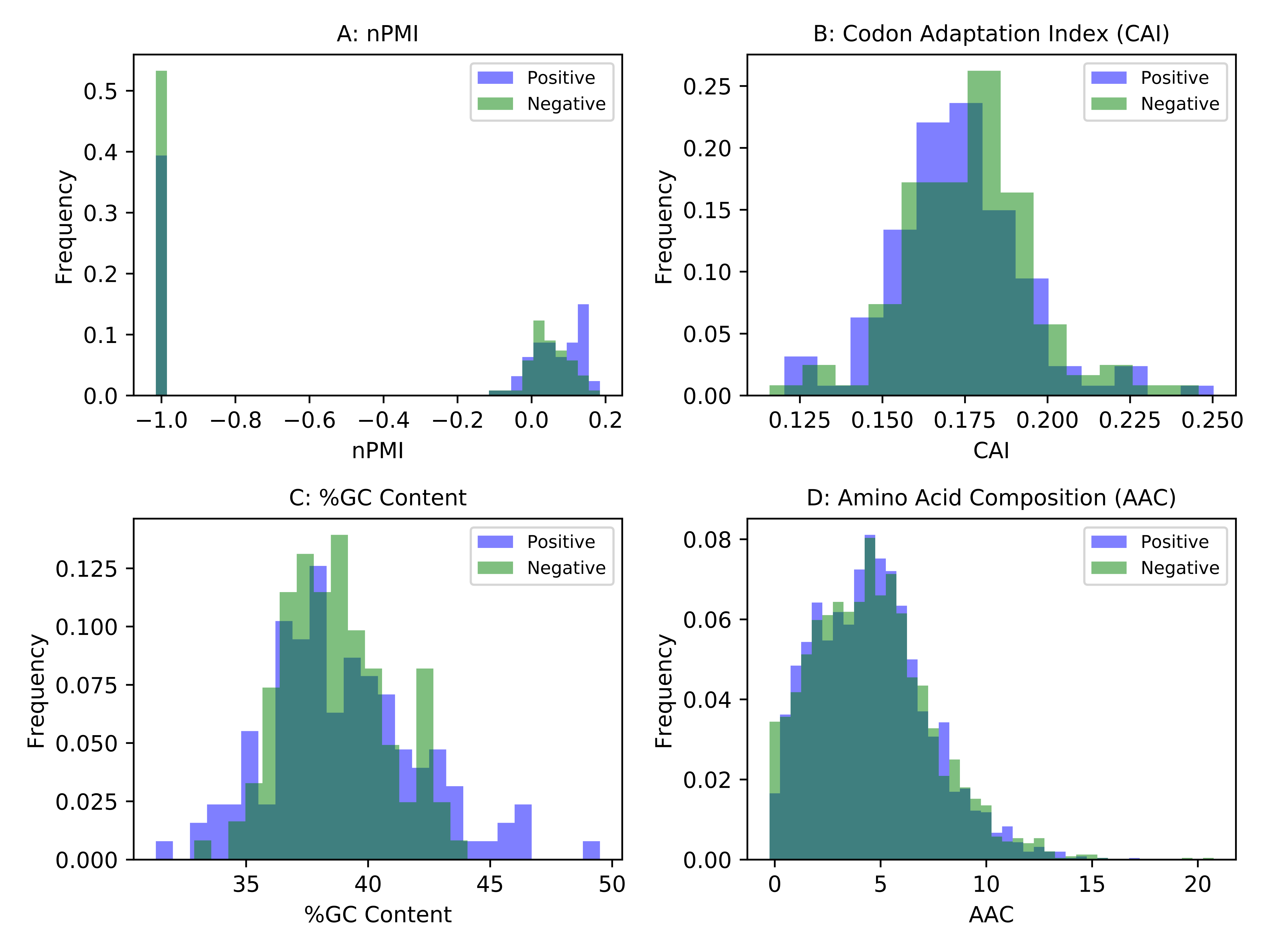

### F4.png

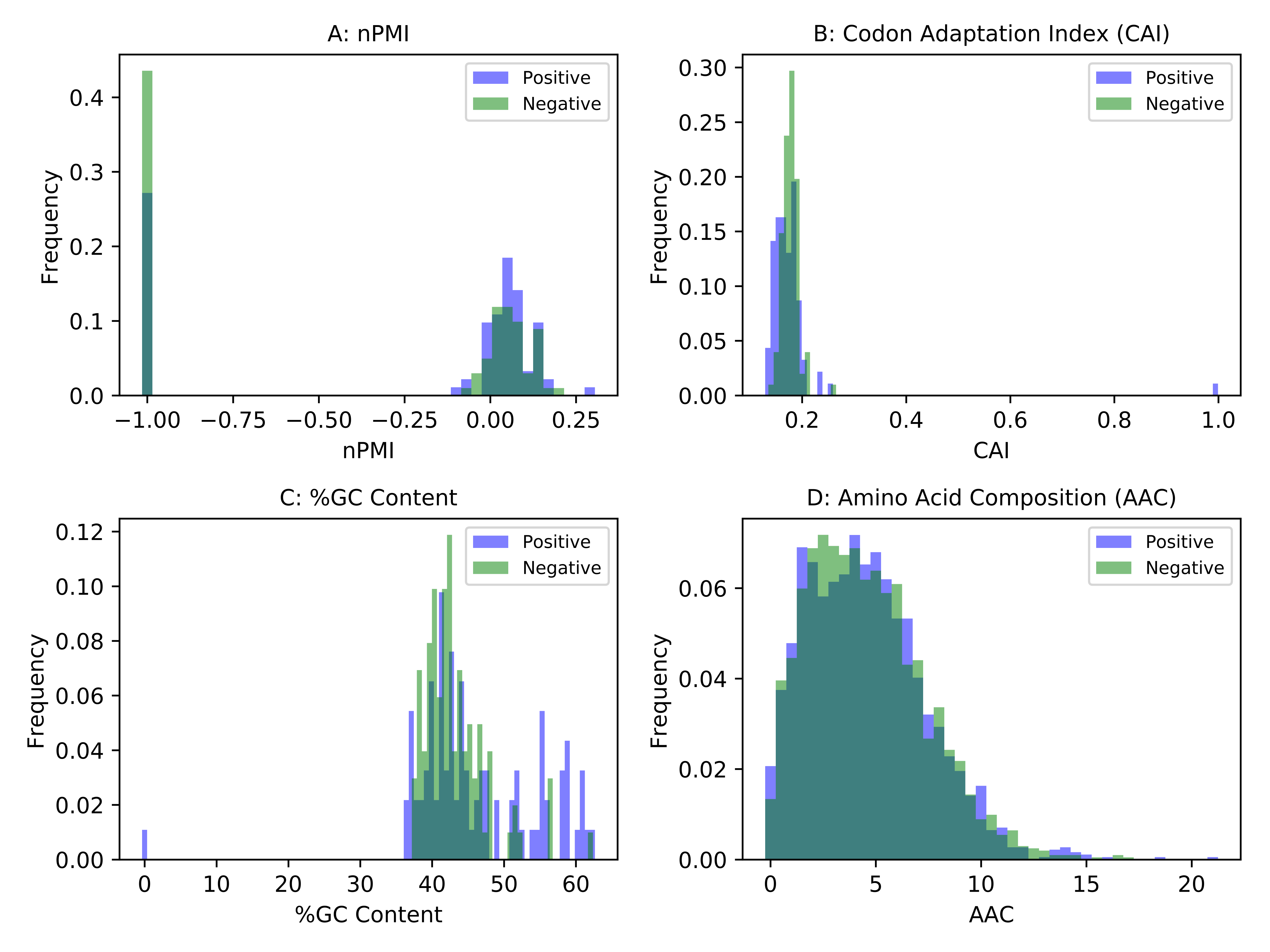

### resamp_ath_50.tiff

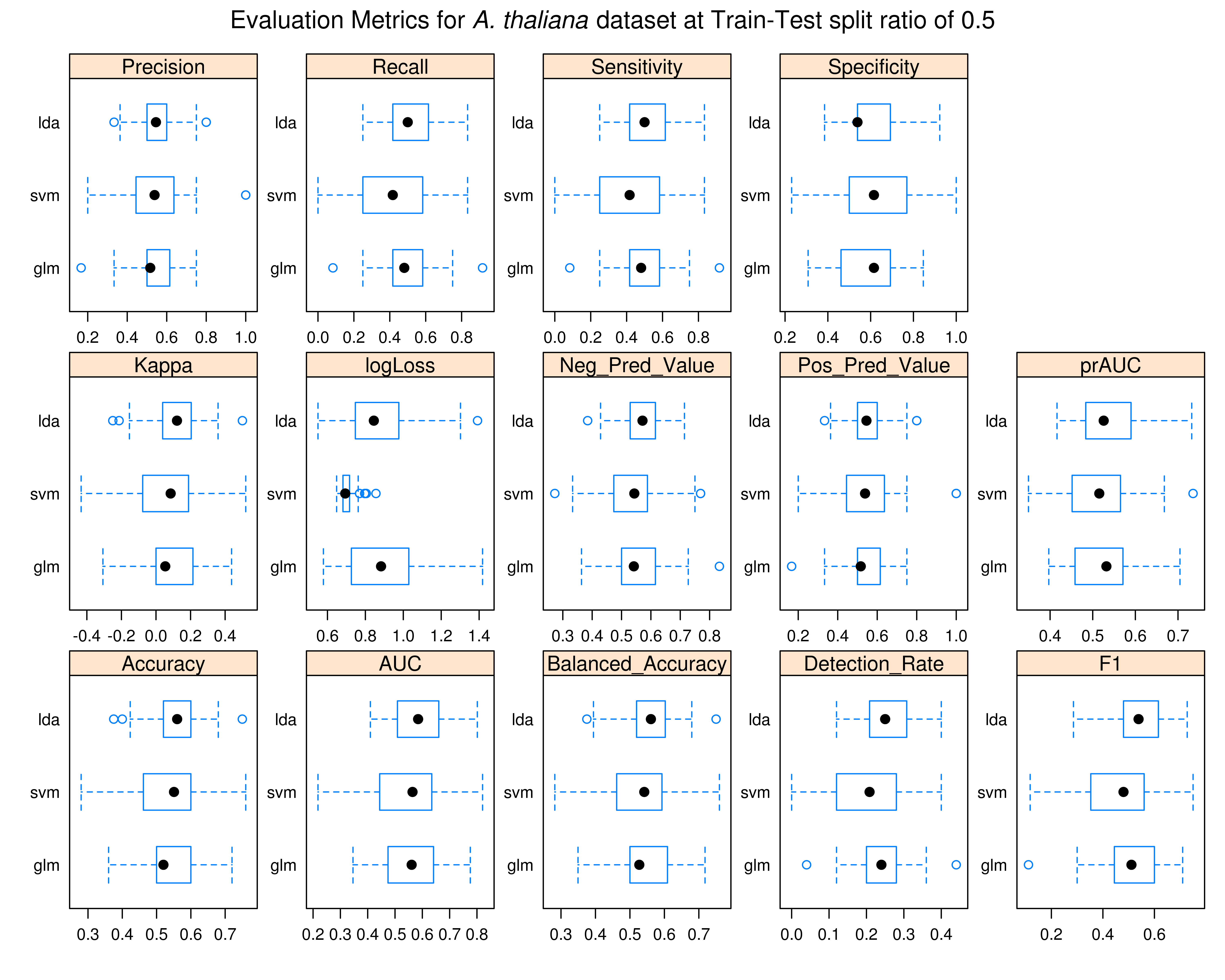

### resamp_ath_60.tiff

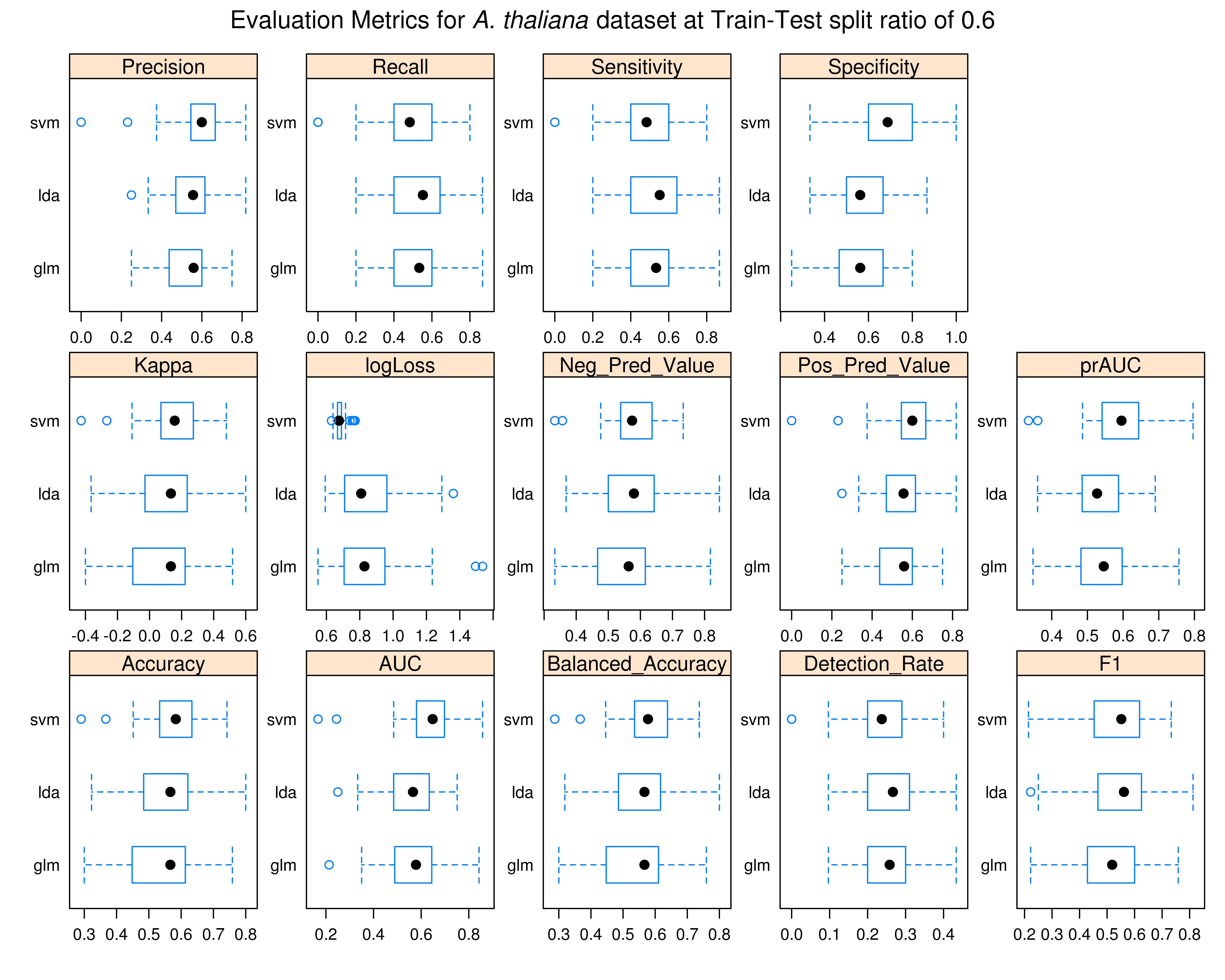

### resamp_ath_70.tiff

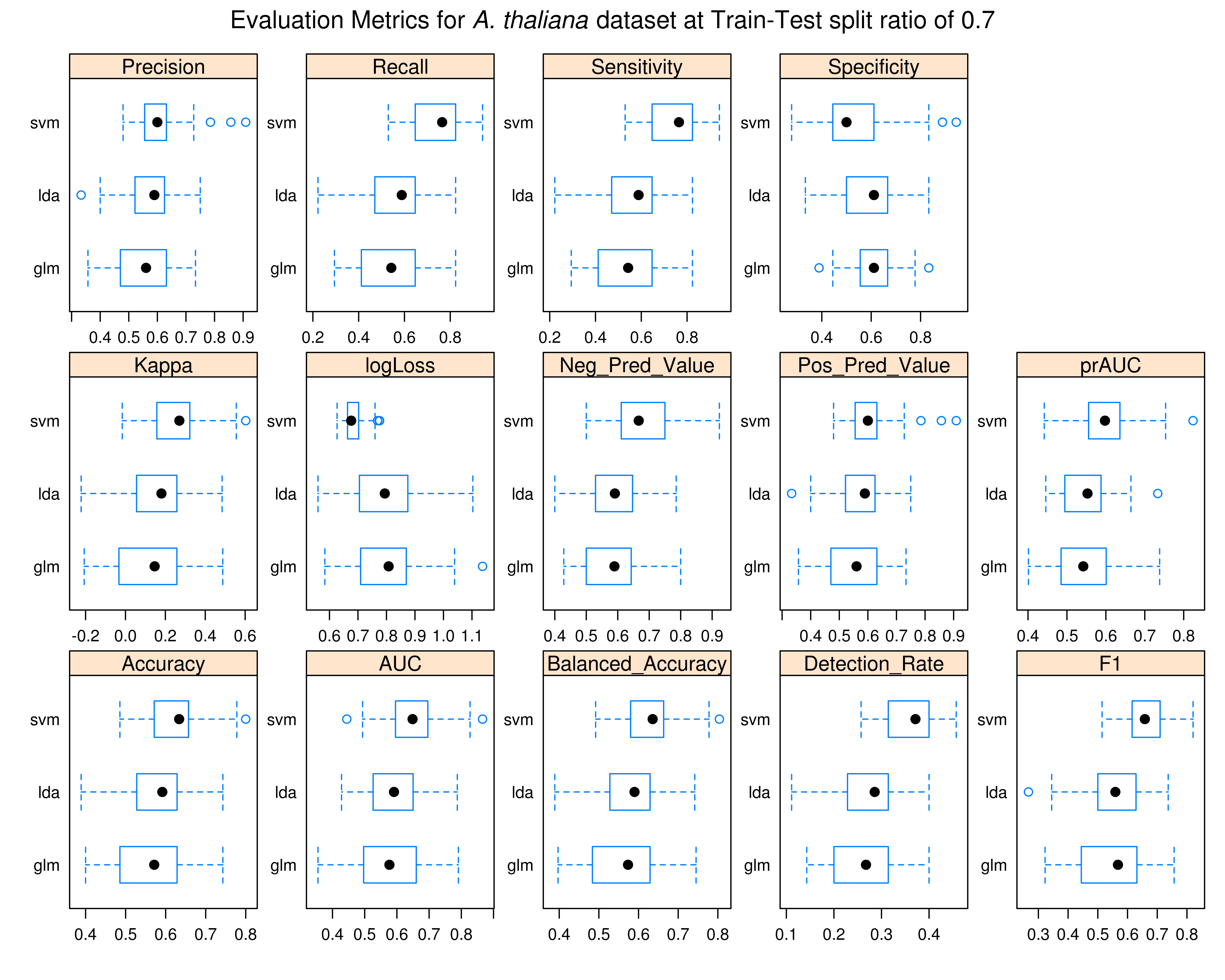

### resamp_ath_80.tiff

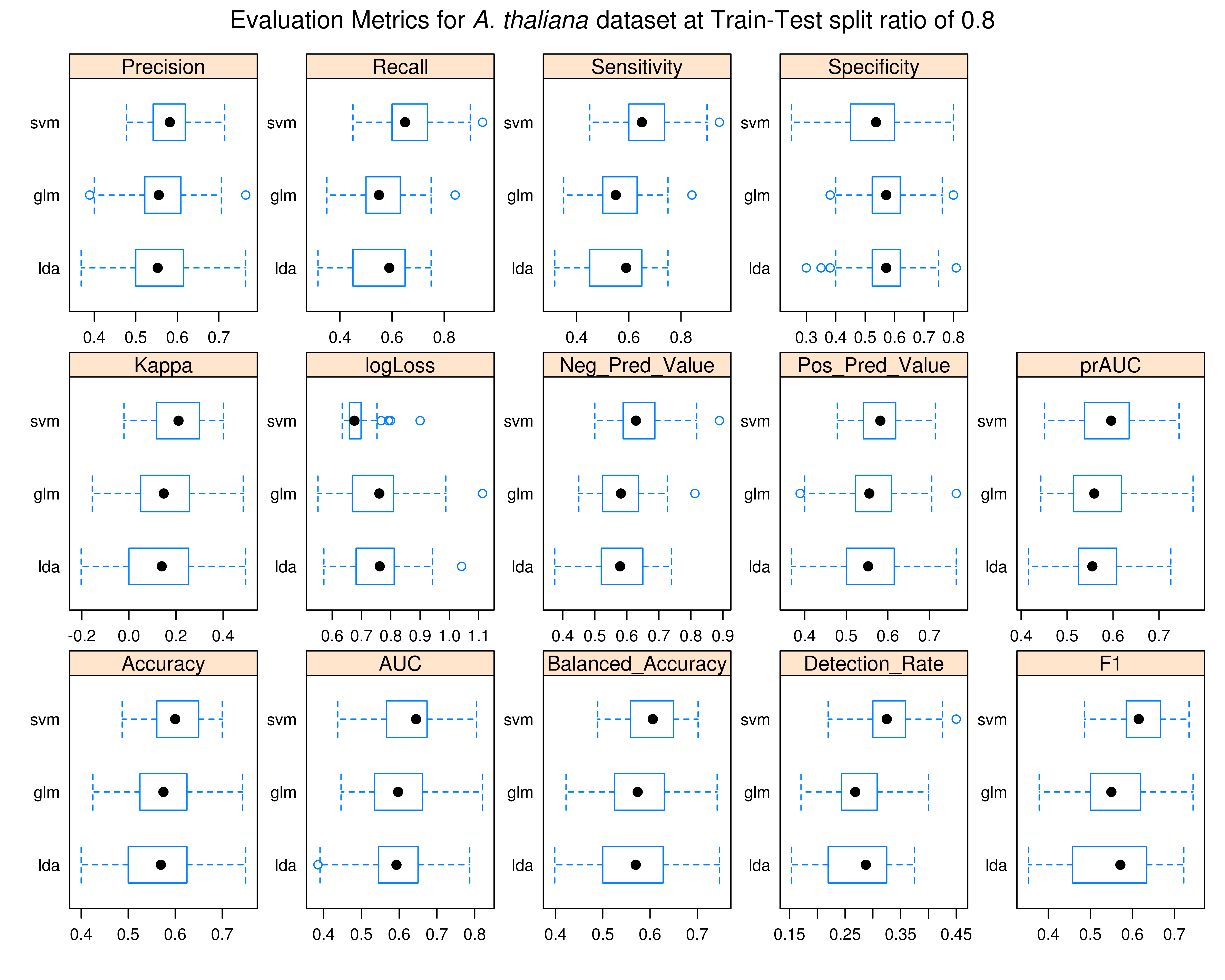

### resamp_ath_90.tiff

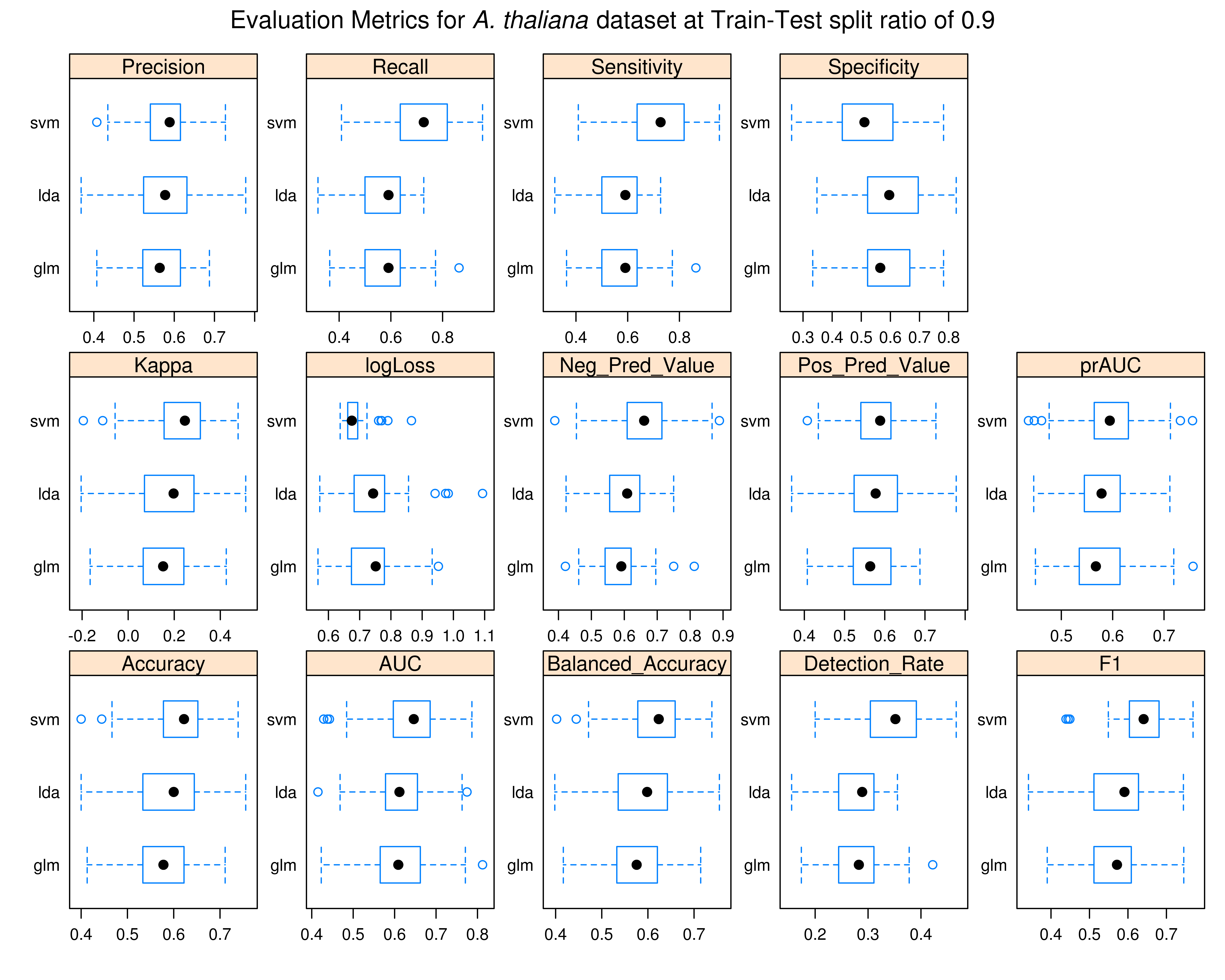

### resamp_coli_50.tiff

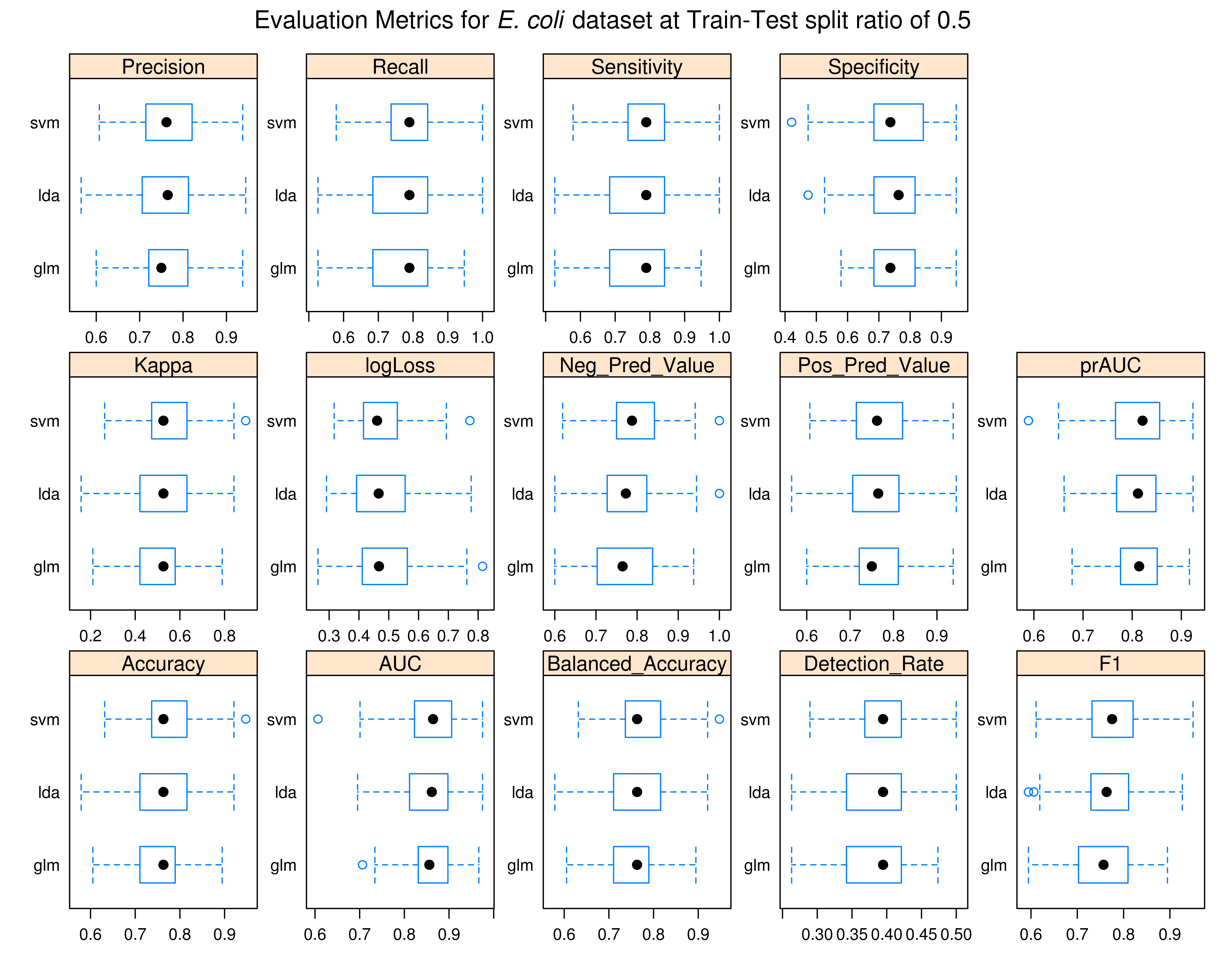

### resamp_coli_60.tiff

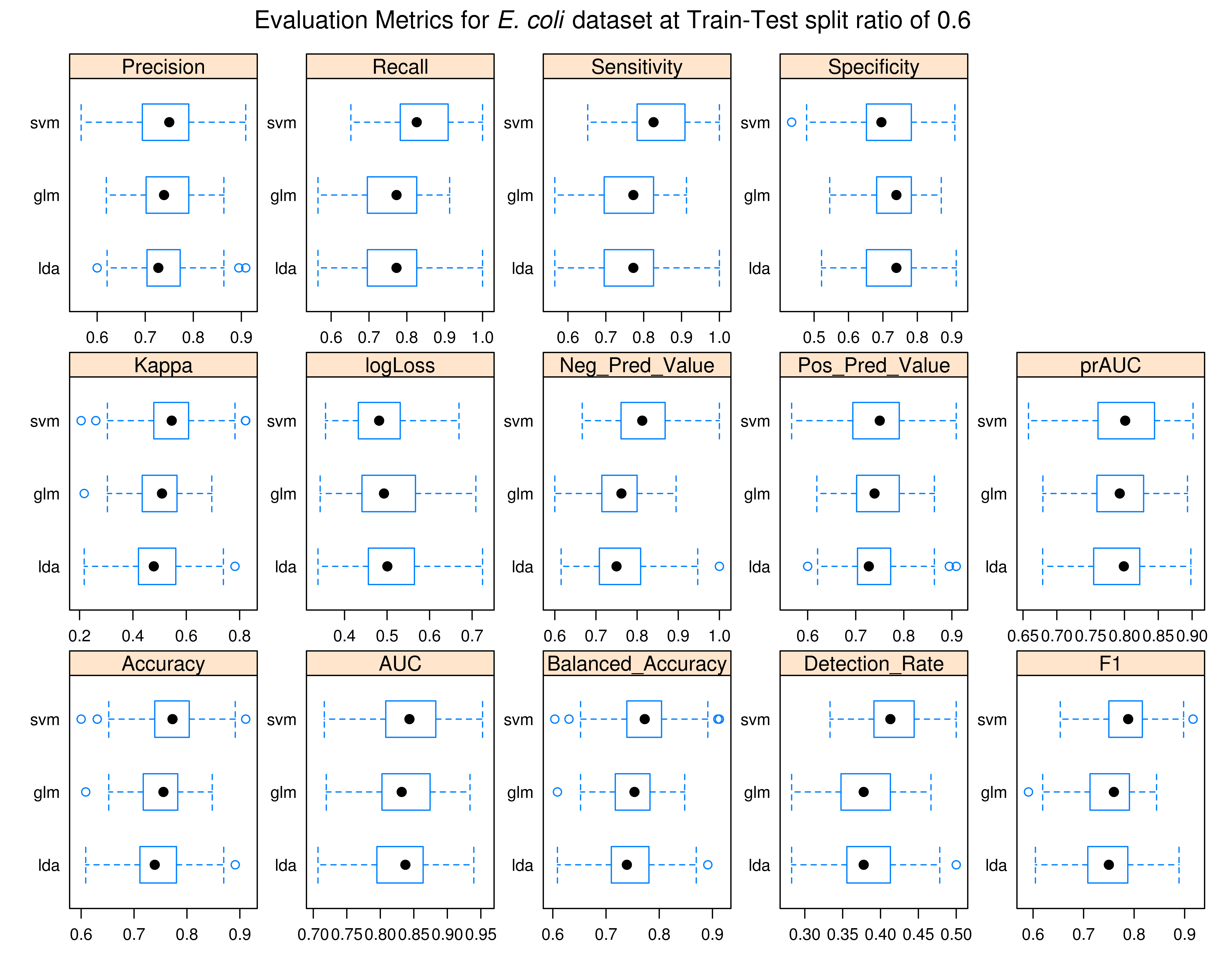

### resamp_coli_70.tiff

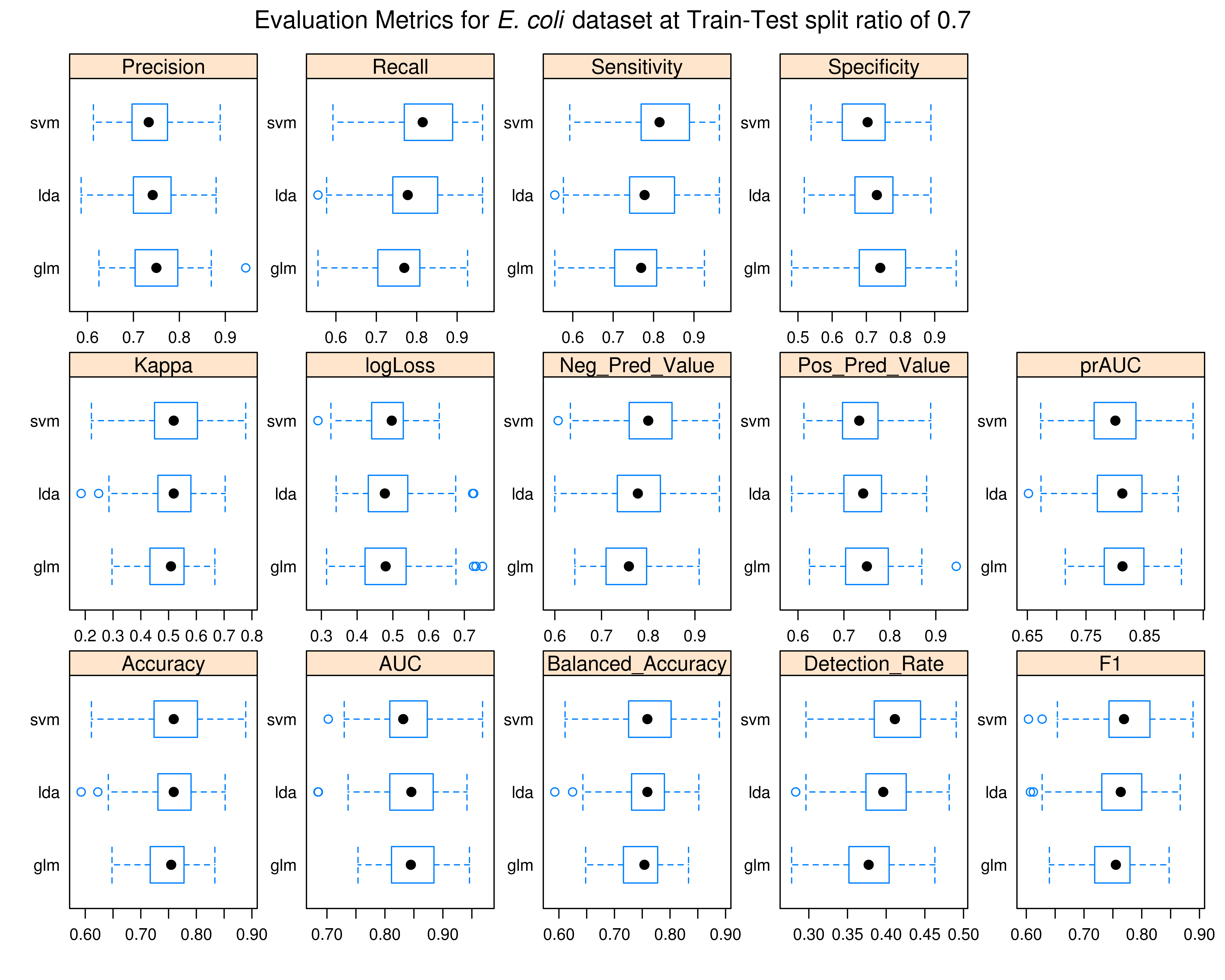

### resamp_coli_80.tiff

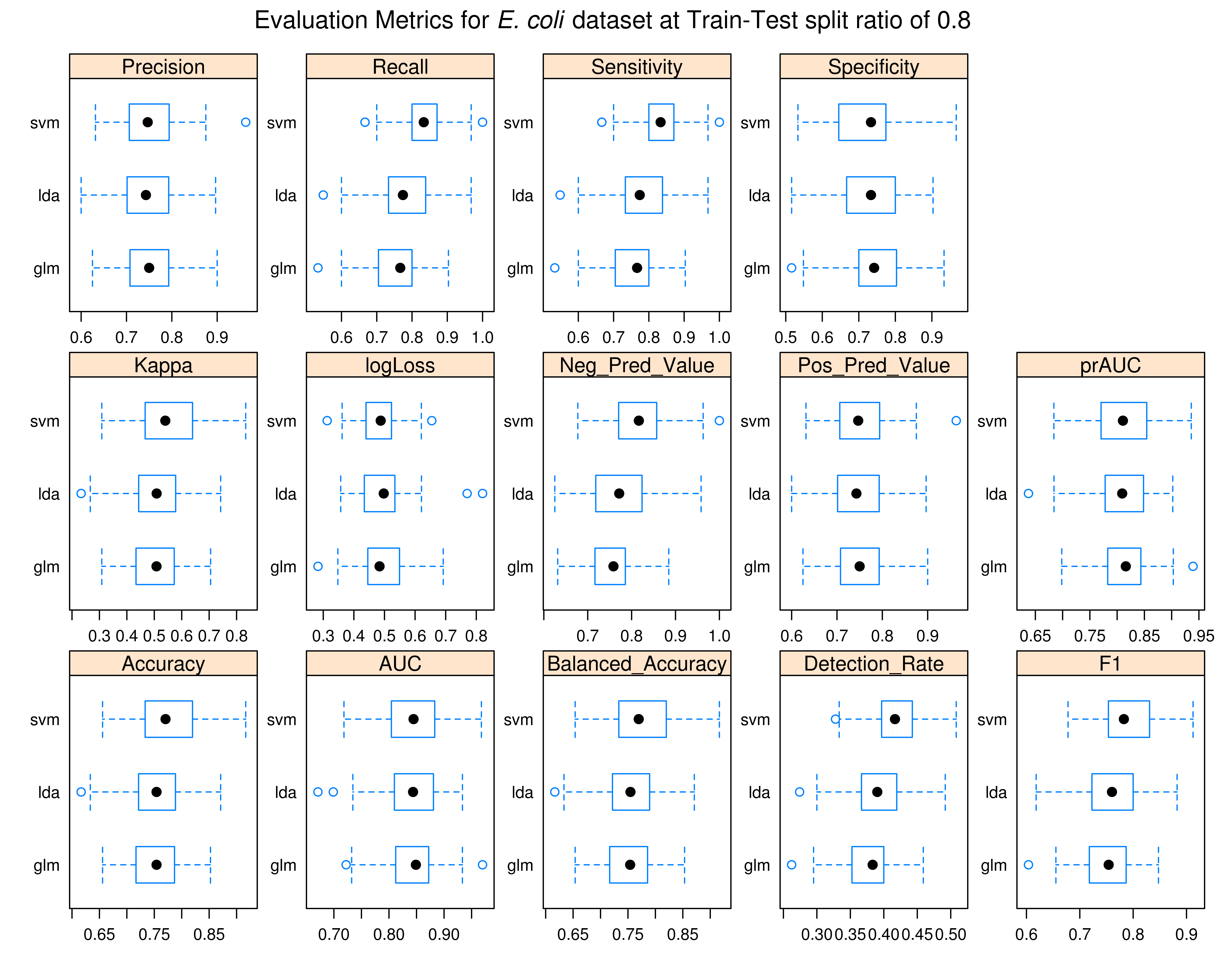

### resamp_coli_90.tiff

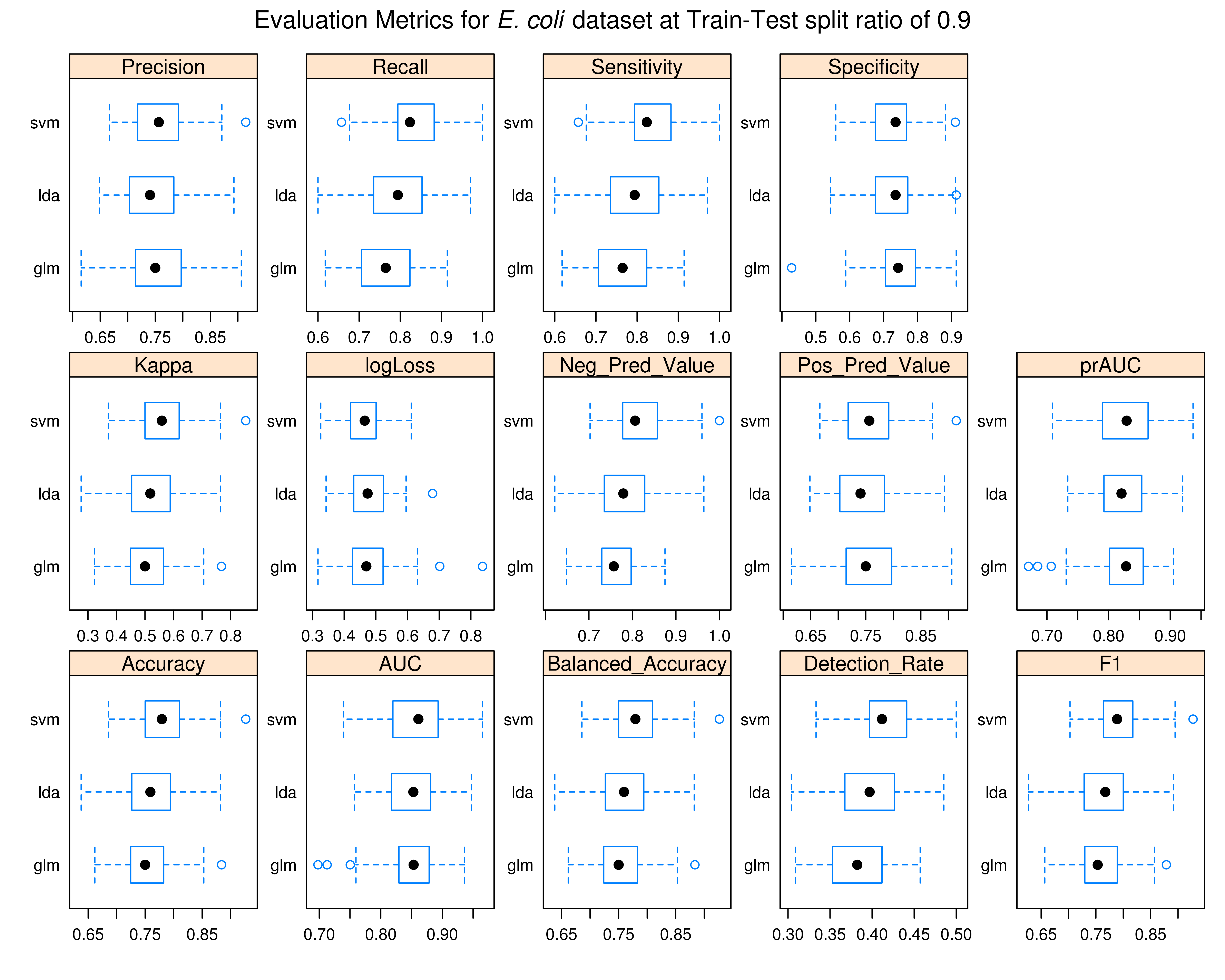

### resamp_osa_50.tiff

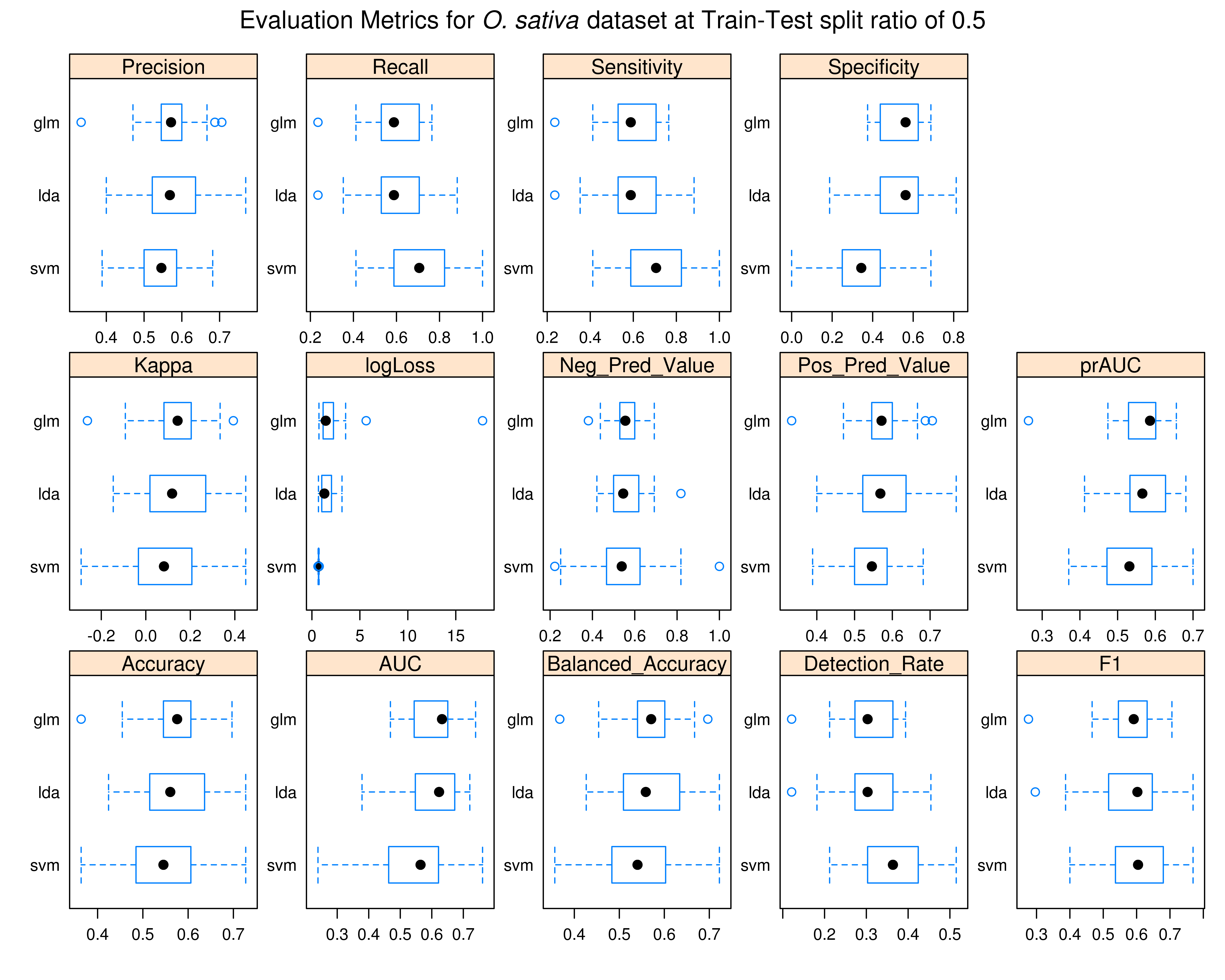

### resamp_osa_60.tiff

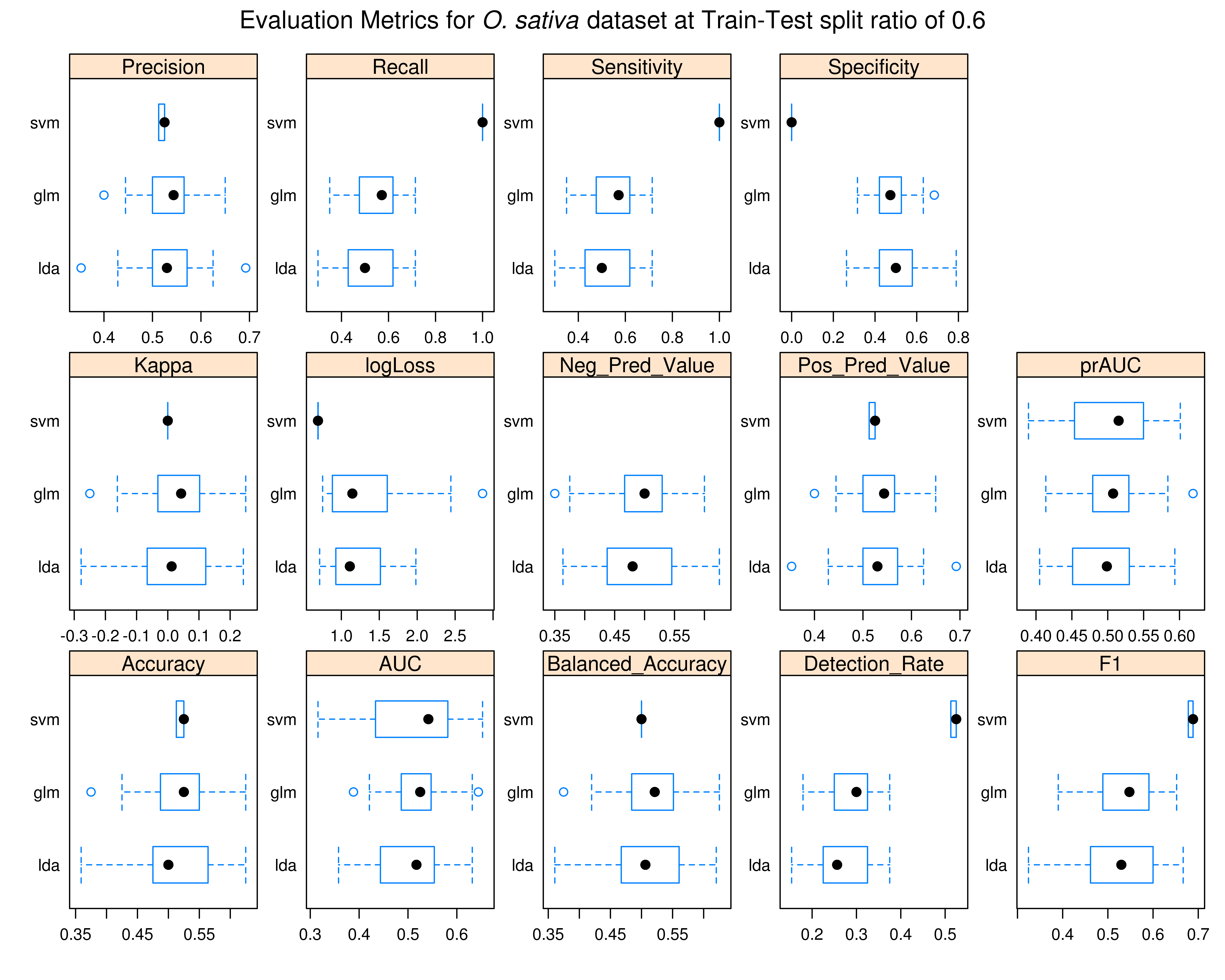

### resamp_osa_70.tiff

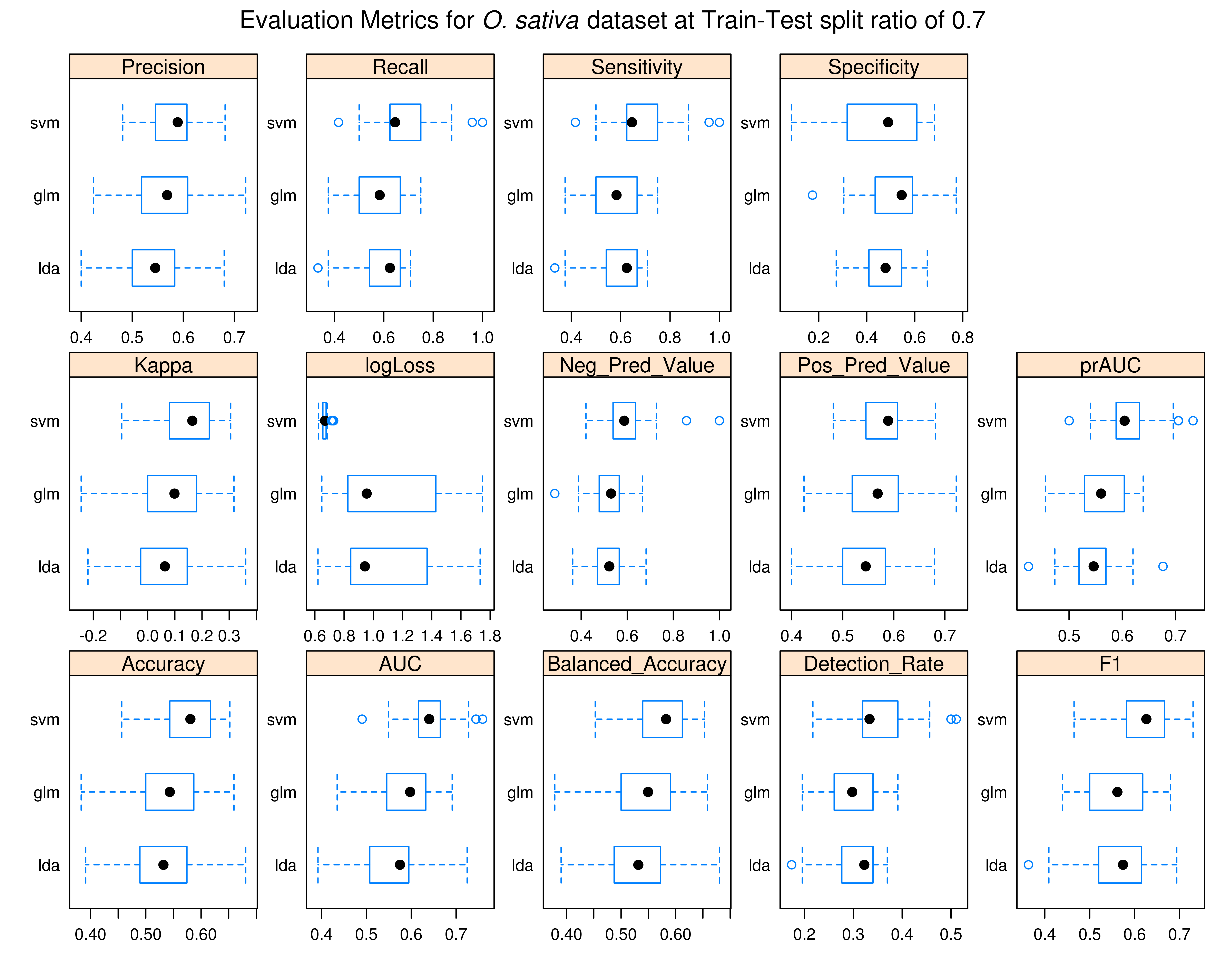

### resamp_osa_80.tiff

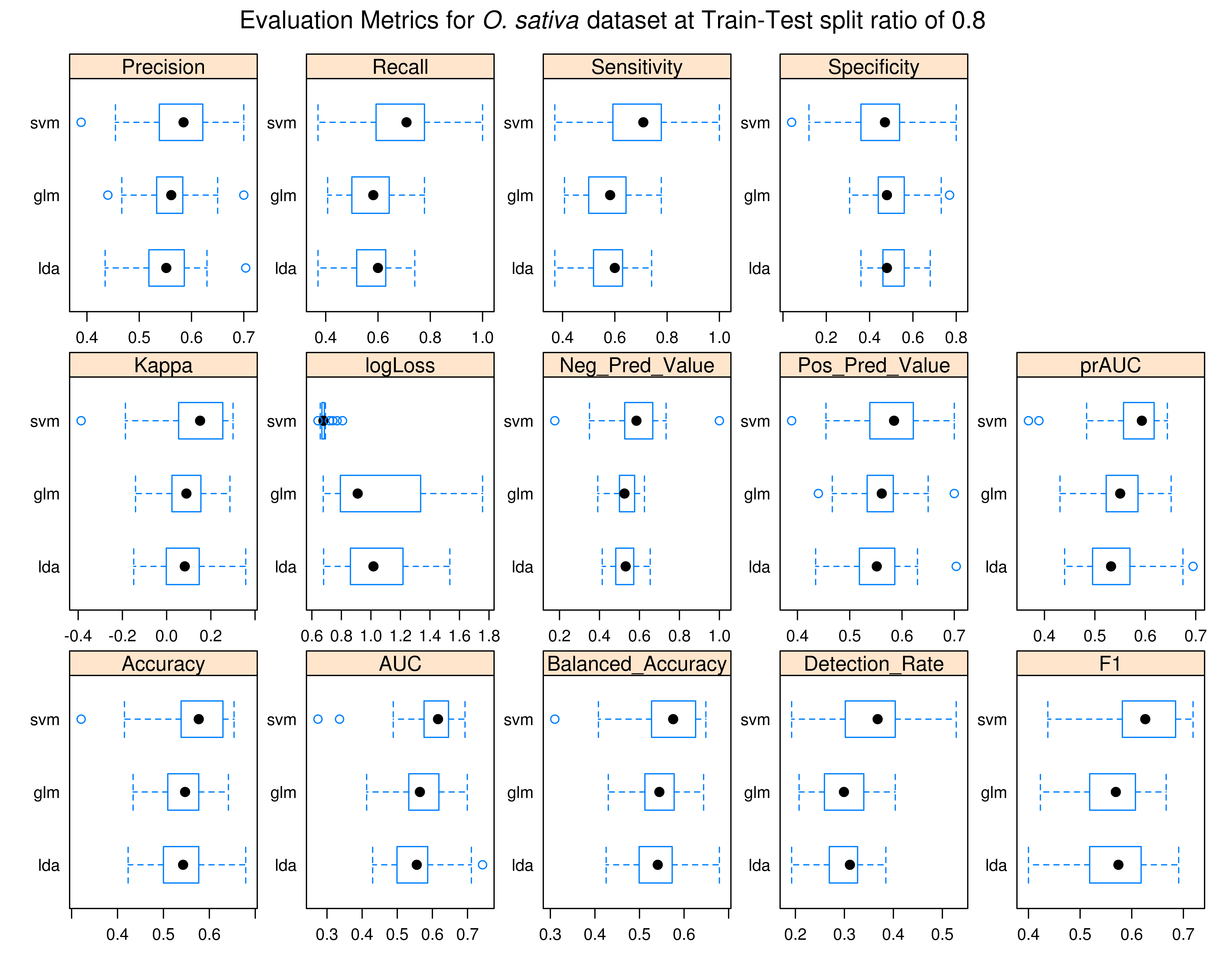

### resamp_osa_90.tiff

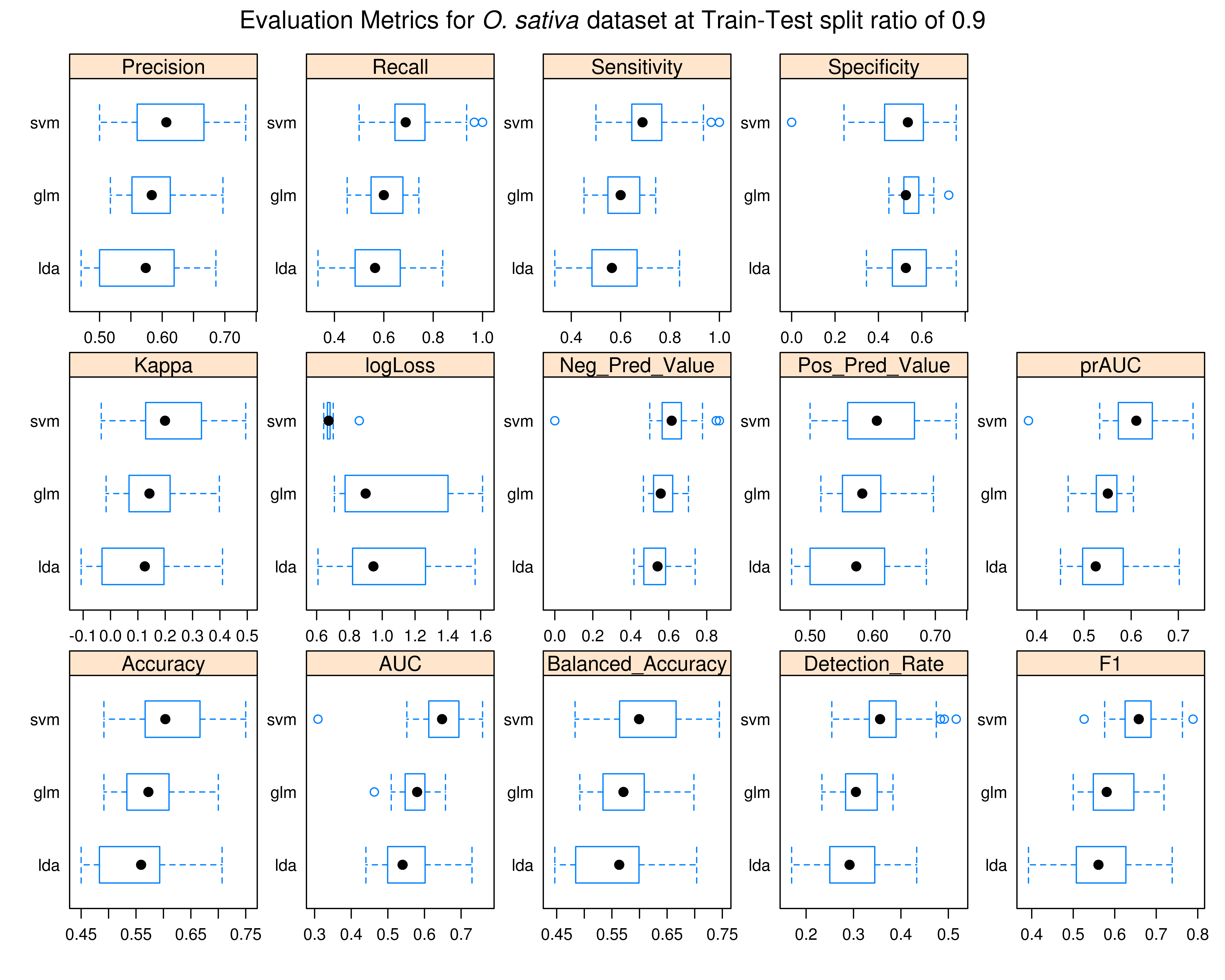

### resamp_sal_50.tiff

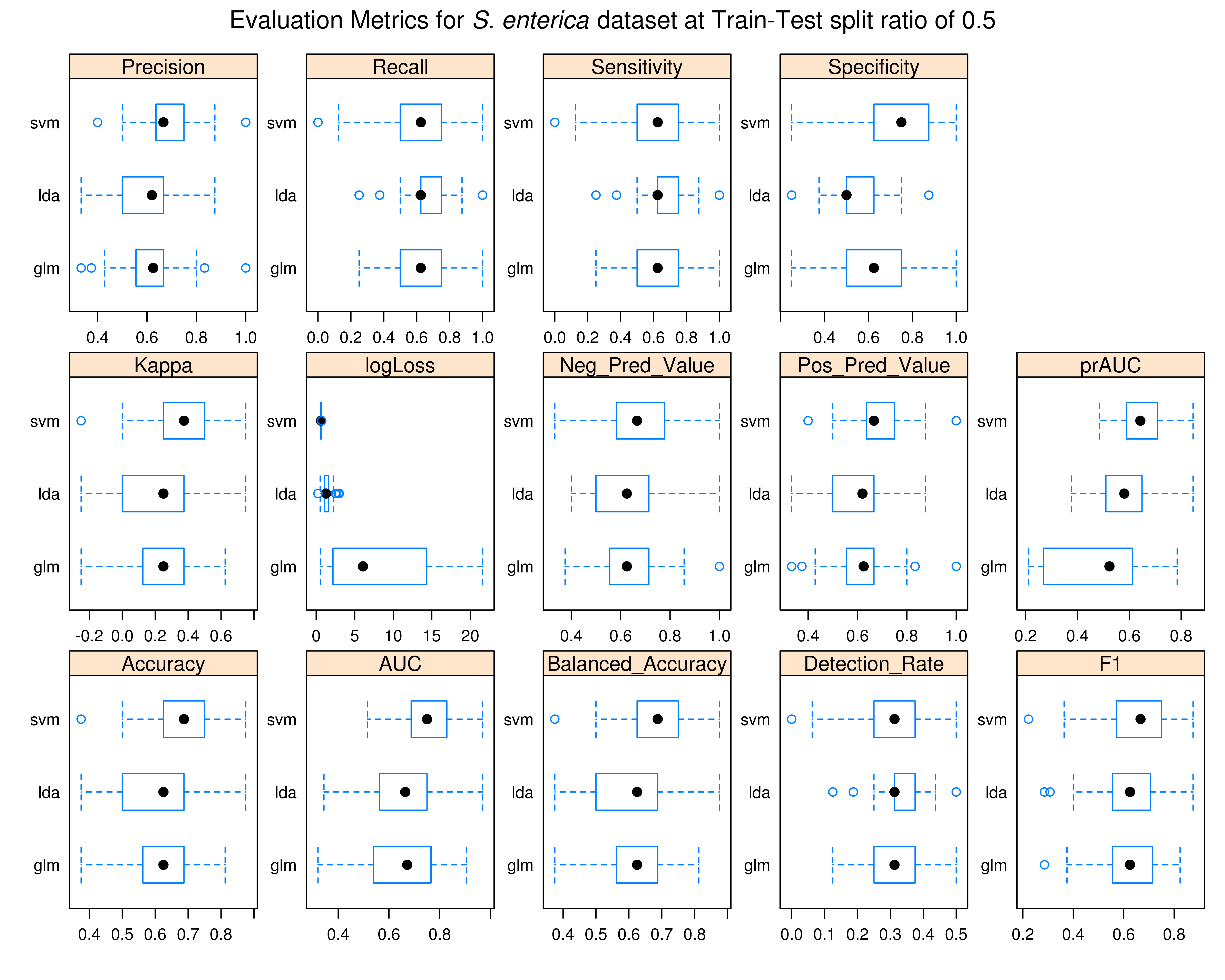

### resamp_sal_60.tiff

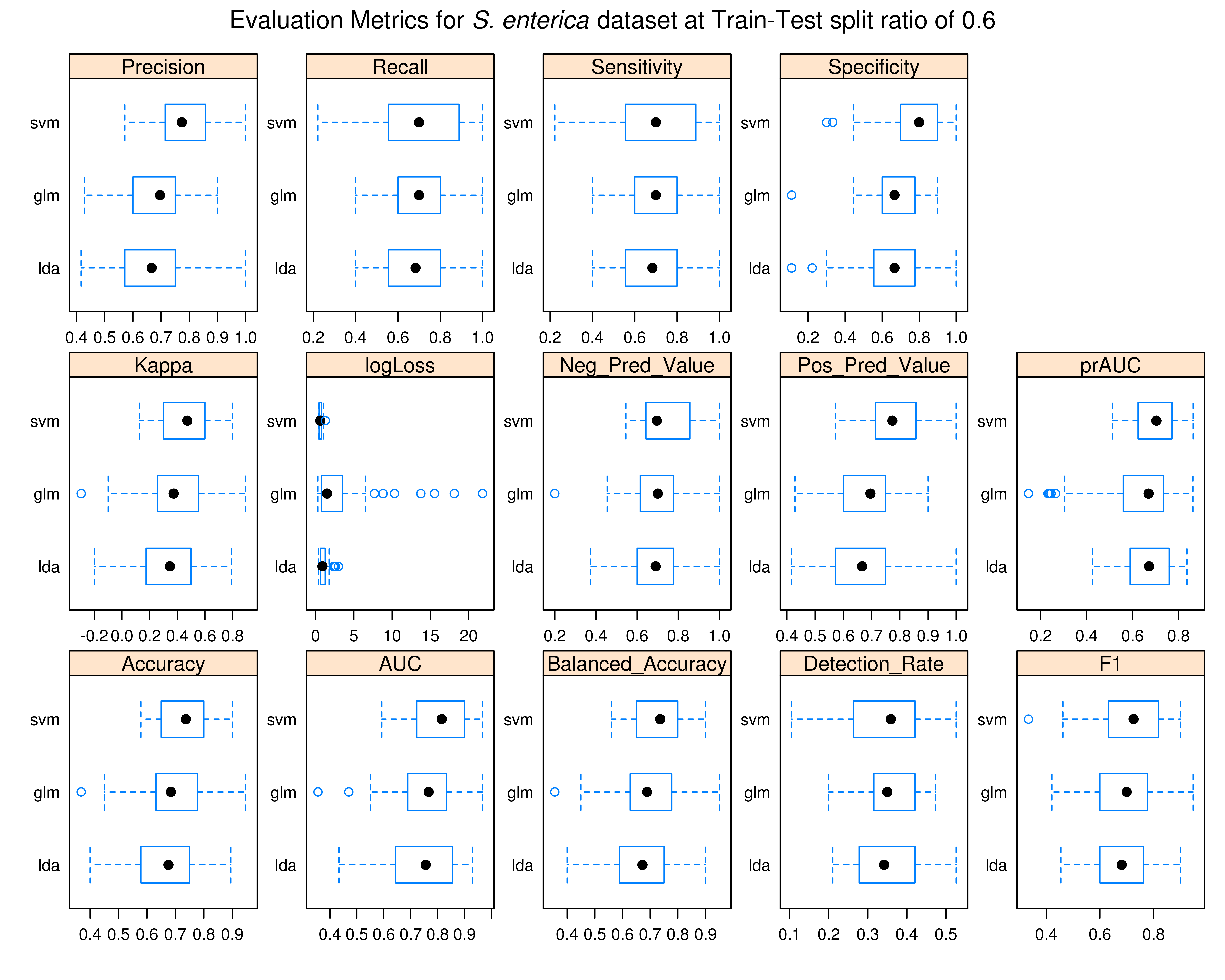

### resamp_sal_70.tiff

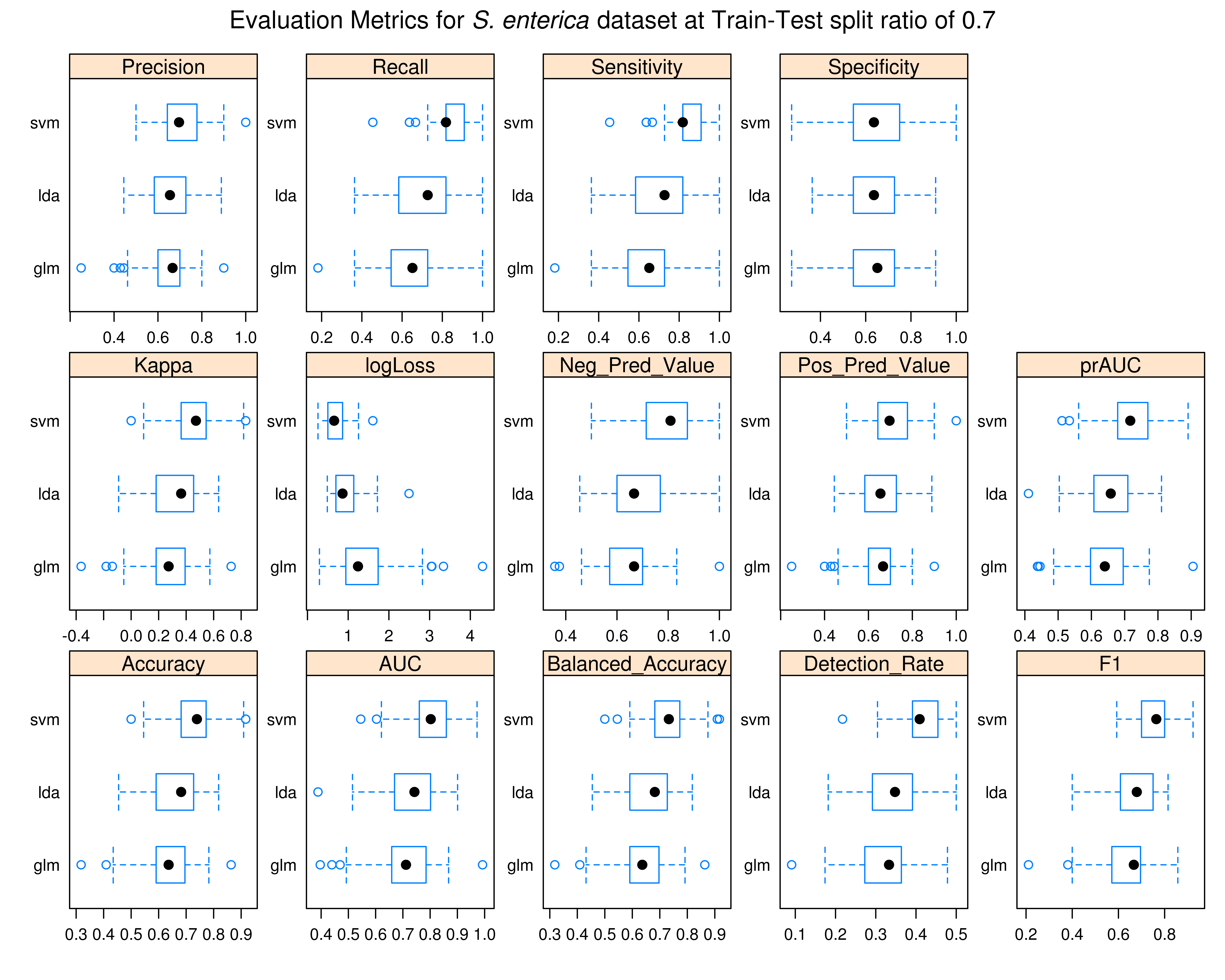

### resamp_sal_80.tiff

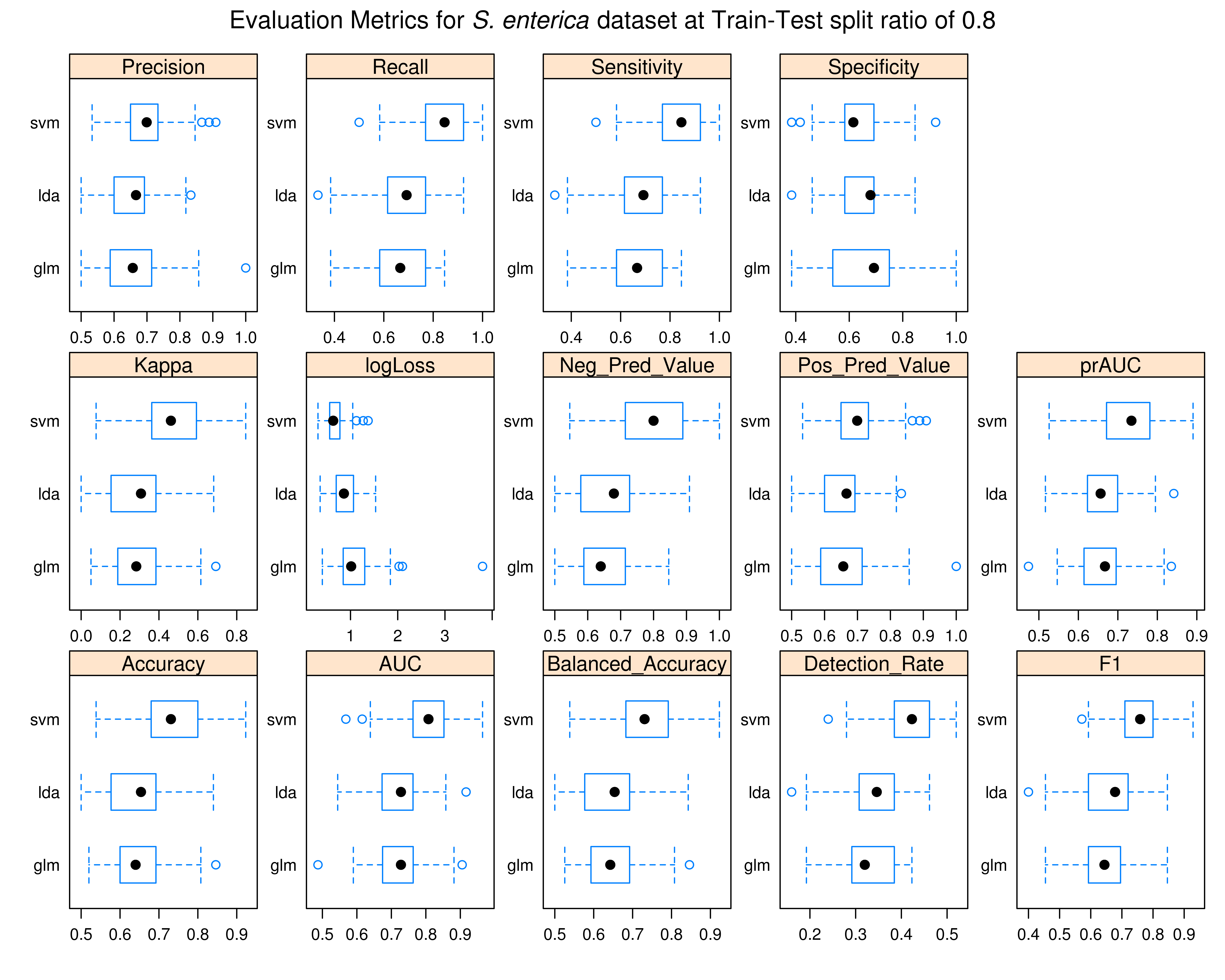

### resamp_sal_90.tiff

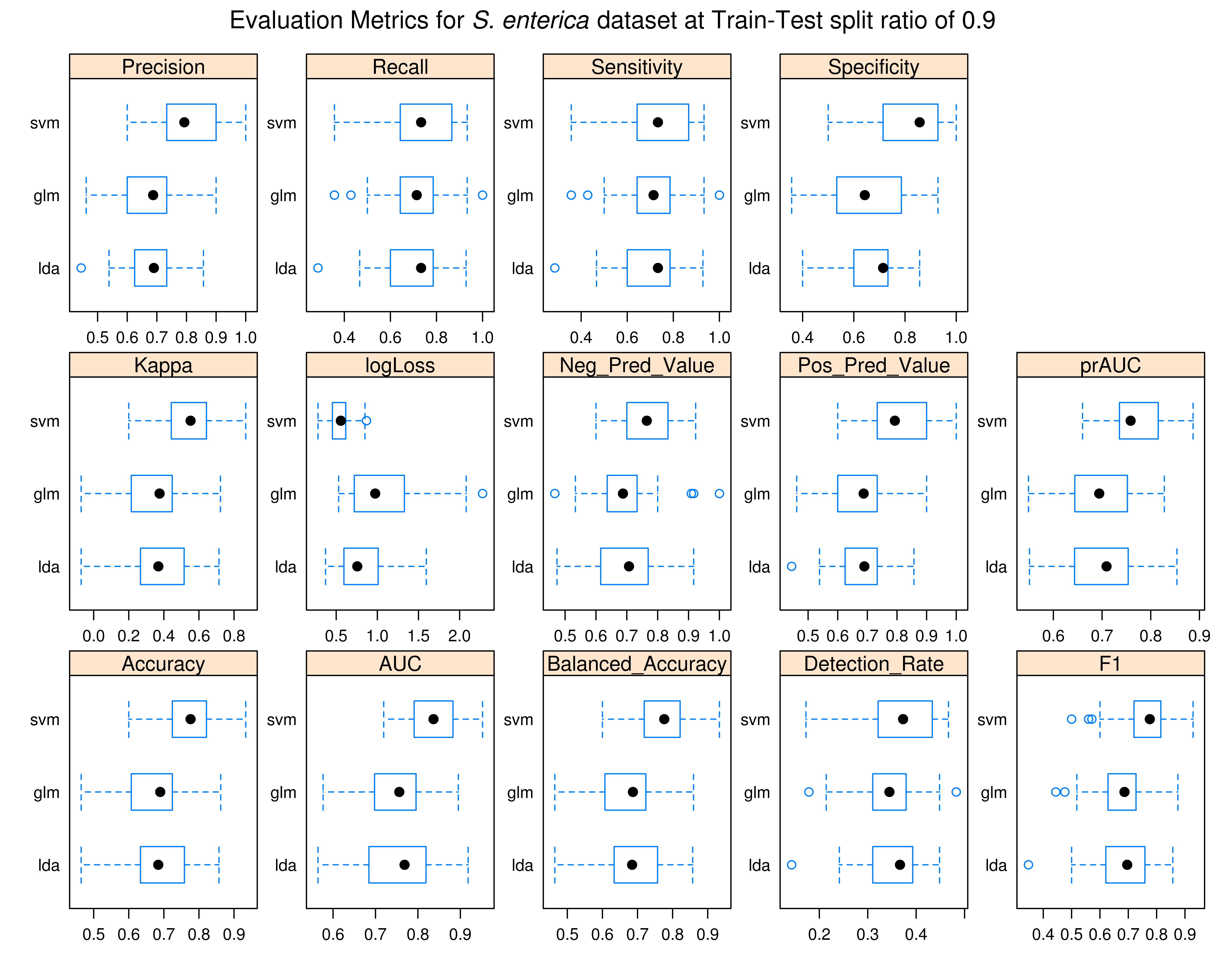
